# Evolution of enzyme levels in metabolic pathways: A theoretical approach

**DOI:** 10.1101/2021.05.04.442631

**Authors:** Charlotte Coton, Grégoire Talbot, Maud Le Louarn, Christine Dillmann, Dominique de Vienne

**Affiliations:** Université Paris-Saclay, INRAE, CNRS, AgroParisTech, GQE – Le Moulon, 91190, Gif-sur-Yvette, France

**Keywords:** Adaptive landscape, Metabolism, Enzyme concentration, Co-regulation, Evolutionary equilibrium, Selective neutrality

## Abstract

The central role of metabolism in cell functioning and adaptation has given rise to countless studies on the evolution of enzyme-coding genes and network topology. However, very few studies have addressed the question of how enzyme concentrations change in response to positive selective pressure on the flux, considered a proxy of fitness. In particular, the way cellular constraints, such as resource limitations and co-regulation, affect the adaptive landscape of a pathway under selection has never been analyzed theoretically. To fill this gap, we developed a model of the evolution of enzyme concentrations that combines metabolic control theory and an adaptive dynamics approach, and integrates possible dependencies between enzyme concentrations. We determined the evolutionary equilibria of enzyme concentrations and their range of neutral variation, and showed that they differ with the properties of the enzymes, the constraints applied to the system and the initial enzyme concentrations. Simulations of long-term evolution confirmed all analytical and numerical predictions, even though we relaxed the simplifying assumptions used in the analytical treatment.

## 1 Introduction

Metabolism is a property of living systems that is central to cell functioning and physiological and genetic adaptation (Cornish-Bowden and Cárdenas 2020). The metabolic networks we see today, comprising an extraordinary diversity of enzymes, cofactors and metabolites, are the product of billion years of evolution that have shaped network topology and enzyme properties.

The importance of metabolism has inspired a wealth of studies on its evolutionary dynamics over different timescales, which follow three main lines of research.

The first line of research relates to the evolution of network structure. Over evolutionary time, continuous selective pressures in diverse environments have resulted in the successive recruitment of new metabolites and enzymes that then coevolve (Noda-Garcia et al. 2018), have modified substrate specificity (Newton et al. 2018) and have determined the general features of the networks, namely functional redundancy, modularity, flux coupling and exaptation (Sambamoorthy et al. 2019). Adaptation to a new environment may involve gain/loss-of-functions and horizontal gene transfers, which rewire the metabolic networks (e.g. Pál et al. 2005; Lee and Palsson 2010; D’Souza and Kost 2016; Morrison and Badyaev 2017).

The second line of research investigates the factors that determine the degree of polymorphism and the rate of evolution of enzyme-coding genes. In addition to classical factors, such as gene dispensability, level of gene expression, protein length, protein folding stability, intron number and protein-protein interaction (see *e.g.* review by Larracuente et al. 2008), various systemic constraints also influence the degree of protein polymorphism and evolution (Pál et al. 2006). Enzymes at branching points of the network or that are highly connected, and thus have pleiotropic effects, are assumed to be more subject to purifying selection; as a consequence, their coding genes have low levels of polymorphism or slow evolutionary rates (Rausher et al. 1999; Vitkup et al. 2006; Greenberg et al. 2008; Rausher 2013). Flux levels and flux control coefficients are additional factors that influence protein evolution: the genes encoding high-flux enzymes have lower evolutionary rates (Colombo et al. 2014), possibly related to enzyme abundance (Aguilar-Rodriguez and Wagner 2018); higher flux control coefficients of enzymes that are upstream in the pathway result in stronger purifying or positive selection (Flowers et al. 2007; Wright and Rausher 2010; Sellis and Longo 2015).

The third line of research is the study of the genetic variability of enzyme properties. The response of organismal fitness to selection pressures depends on genetically variable enzyme properties, namely kinetic parameters, thermodynamic stability and concentration. In the context of the historical debate between neutralism and selectionism, a number of emblematic examples of variation in enzyme properties have been studied in detail. For instance, in *Drosophila*, the fast and slow allozymes of alcohol dehydrogenase differ both in the value of their catalytic constant (the turnover number, *k*_cat_) and their abundance; also in *Drosophila* the A and B variants of G6PD have different quaternary structures, thermal stability and *K*_m_ values; in *Fundulus heteroclitus*, the allozymes of LDH-B display kinetic, abundance and stability differences, etc. (reviewed in Eanes 1999). Furthermore, quantitative proteomic studies in various species have revealed the breadth of the genetic variability, the polygenic nature and the relationship with integrated phenotypic traits of enzyme concentrations (Damerval et al. 1994; Blein-Nicolas et al. 2013; Albert and Kruglyak 2015; Chick et al. 2016; Jiang et al. 2019).

In this context, the evolution of enzyme properties in response to selective pressures on the flux, considered to be a proxy for fitness, is an important issue. This question has been addressed in relation to protein folding stability, catalytic constant values and enzyme concentration, but to our knowledge the number of studies is quite low. Mutations that increase protein folding stability are favored by selection, with evolution towards near neutrality up to the mutation–selection balance steady state (Serohijos and Shakhnovich 2014; Bershtein et al. 2017). Regarding enzyme activity, Kacser and Beeby (1984), using a systems biology approach, proposed a model of early enzyme evolution in which selection for faster growth causes a transition from few, multifunctional, poorly active enzymes to many, differentiated, monofunctional enzymes with high *k*_cat_ values. However, Heckmann et al. (2018) recently showed that the systemic properties of a network result in epistatic interactions that maintain *k*_cat_ values far from their theoretical maximum, preventing fitness from reaching its optimum. This result can be explained by the relationship of diminishing returns between turnover number and flux: the effect of a given increase in *k*_cat_ on the flux vanishes as *k*_cat_ increases, so that the flux eventually reaches a plateau.

Interestingly, the same type of saturation curve was invoked by Hartl et al. (1985) to account for the widespread selective neutrality of enzyme polymorphisms. These authors analyzed the evolution of enzyme quantity, equated to activity, under selection for higher flux, and showed that successive mutations affecting a particular enzyme drive the enzyme concentration towards the plateau of the concave flux-enzyme curve, where the mutations do not have any detectable effect, *i.e.* they are selectively nearly neutral. This geometrical model of the “natural selection of selective neutrality” is attractive, but surprisingly it has not been assessed with an adaptive dynamics approach. In addition, this model considered only a single enzyme in a given pathway and imposed no upper limits on enzyme quantities, even though there are well-known constraints on space and energy in the cell (Koehn 1991; Kurland and Dong 1996; de Vienne et al. 2001b; Ellis 2001; Eguchi et al. 2018; Klumpp et al. 2019). Although these points were raised, the evolutionary implications were not analyzed by Hartl et al. (1985). The question of the optimization of enzyme kinetic parameters and concentrations in metabolic pathways was analyzed by Klipp and Heinrich (1994, 1999). However, their approach did not include any evolutionary genetics concept and did not allow them to grasp the dynamics of the evolutionary process.

Here, we develop the first adaptive dynamics model of the evolution of enzyme concentrations when there is directional selection for increased flux in a pathway. Our model includes two biologically realistic features: (i) the concentration of each enzyme in the pathway can be altered by mutation, and (ii) constraints, specifically competition for cellular resources and/or co-regulation, can prevent enzyme concentrations from varying freely. To formalize the relationship between enzyme concentrations and flux, we used the metabolic control theory (MCT) framework (Kacser and Burns 1973; Heinrich and Rapoport 1974; Kacser et al. 1995), as generalized by Heinrich et al. (1991) and Lion et al. (2004) to take into account the possible interdependence of enzyme concentrations.

First, we present a system of ordinary differential equations that describes the evolution of enzyme concentrations in a pathway. We then show how the flux evolves and how it determines the evolutionary equilibria of the relative enzyme concentrations. We demonstrate analytically, and validate using computer simulations, that a theoretical equilibrium of relative enzyme concentrations exists whatever the conditions, but that this equilibrium is not necessarily reached when the flux is limited by constraints on enzyme concentration variation. Thus, we differentiate the *effective* equilibrium, *i.e.* the equilibrium that is actually reached, from the *theoretical* equilibrium. When there is co-regulation in addition to competition, the initial enzyme concentrations affect the evolutionary outcome. Finally, we show that multi-enzyme selective neutrality, *i.e.* the selective neutrality of all enzymes, is achieved whatever the constraints, but that the size of the range of neutral variation largely depends on constraint-specific parameters and possibly on the initial state of the system.

## 2 Material and Methods

### 2.1 Theoretical developments

#### 2.1.1 The metabolic model

##### 2.1.1.1 Flux expression

We considered a metabolic pathway of *n* unsaturated Michaelian enzymes transforming substrate X_0_ into product X_*n*_ *via* a series of unimolecular reversible reactions:

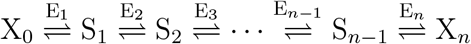

At steady state, the metabolic flux *J* through this pathway is (Kacser and Burns 1981):

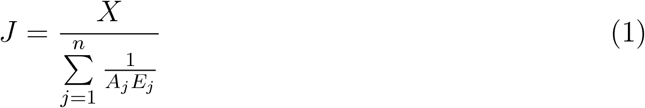

where:

- *X* = *X*_0_ − *X_n_/K*_1,*n*_. *K*_1,*n*_ is the product of the successive equilibrium constants of all reactions. *X* is assumed to be constant and symbolizes that the environment is constant.
- 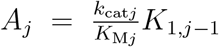, where *k*_cat*j*_ is the turnover number of enzyme *j*, *K*_M*j*_ is the Michaelis-Menten constant of the enzyme *j* for its substrate and *K*_1,*j*−1_ is the product of the successive equilibrium constants up to the (*j* − 1)^th^ reaction: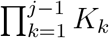. Thus *A_j_* is a composite parameter that includes both catalytic efficiency 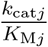 and a product of thermodynamic constants that depends on the enzyme’s position in the pathway. Hereafter, *A_j_* is referred to as the “pseudo-activity” of enzyme *j*.
- *E_j_* is the concentration of enzyme *j*. Note that *E_j_* in italics denotes the concentration of enzyme *j* while E_*j*_ in Roman letter denotes enzyme *j* (see Glossary of mathematical symbols in Appendix A).

Considering the total enzyme concentration, 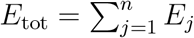, and the relative concentration of enzyme *j*, 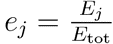, equation 1 becomes:

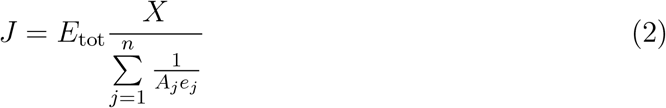

The flux can be modified by mutations affecting 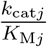 or *E_j_*. We considered only the latter, because the variability of enzyme concentrations is by far the main source of variation in enzyme activity in populations and species *(see Discussion)*.

##### 2.1.1.2 The effects of mutation when enzyme concentrations are interdependent

Let a mutation of small effect *ν* targeting concentration *E_i_* of enzyme *i* (*ν* is positive or negative). We assume that the probability law of mutation effect *ν* is identical for all enzymes, so *ν* is not indexed. The concentration of enzyme *i* after a mutation is:

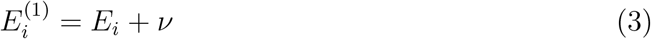

Because of constraints such as resource limitations or complex interactions between cell components, this change may not correspond to the actual effect of the mutation on the concentration of enzyme *i* and/or it may modify the concentration of other enzymes. Considering two causes of interdependence of enzyme concentrations, separately or jointly, we formalized the consequences of a mutation in the following way:

###### 1. Co-regulations

Various direct and indirect molecular mechanisms result in coregulation (either positive or negative) of enzyme concentrations. Thus, mutations can jointly modify the concentrations of co-regulated enzymes *(see Discussion)*. To express formally the effect of a mutation targeting the concentration of enzyme *i* (equation 3) on the concentration of enzyme *j*, we write:

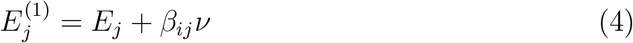

where *β_ij_*, the slope of the relation between *E_i_* and *E_j_*, is the *co-regulation coefficient*. We assume linearity between co-regulated enzymes, so we use constant co-regulation coefficients and we pose *β_ii_* = 1, *β_ij_* = 1*/β_ji_* and, for any triplet of co-regulated enzymes (*i, j, k*), *β_ij_* = *β_ik_β_kj_*. If enzymes *i* and *j* are not co-regulated, we have *β_ij_* = *β_ji_* = 0. All pairwise co-regulation coefficients can be grouped in a co-regulation matrix, noted **M**_*β*_ (Supporting Information SI.B.1.1).

For *n* co-regulated enzymes, the total enzyme concentration after a mutation affecting enzyme *i* is:

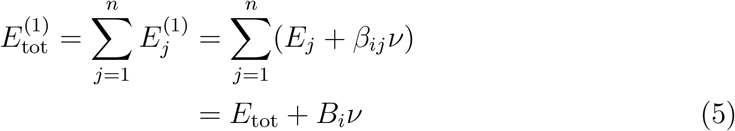

where 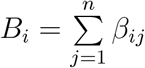 quantifies the impact on *E*_tot_ of a mutation targeting *E_i_*. Indeed, from equations 3 and 5, we have:

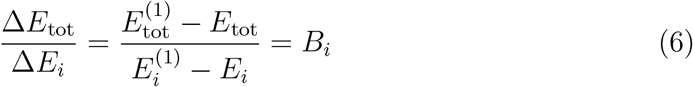

*B_i_* is called the *global co-regulation coefficient* (Supplementary information SI.B.1.2).

###### 2. Competition for resources

Cellular resources are limited in terms of space, energy, the availability of ribosomes, etc. The total enzyme concentration a cell allocates to a given pathway is necessarily bounded, so that an increase (resp. decrease) in the concentration of an enzyme due to a mutation can result in a decrease (resp. increase) in the concentration of other enzymes (Snoep et al. 1995; Albertin et al. 2013). We modeled this trade-off by fixing *E*_tot_. As a consequence, if a mutation of effect *ν* targets enzyme *i*, the concentrations of all the other enzymes will be modified, as will be the effect of the mutation on enzyme *i*. This can be viewed as a two-step process:

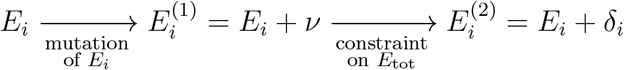

where *δ_i_* is the mutation effect after applying the constraint on total enzyme concentration, which changes from 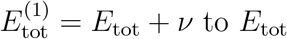. This constraint is assumed to respect the proportionality between concentrations, *i.e.*

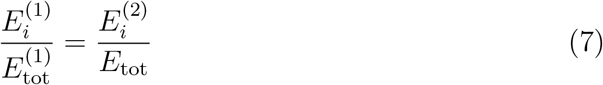

From this relation, the expression of *δ_i_* can be deduced (Supporting Information SI.B.1.3):

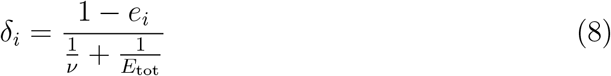

where 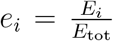. We call *ν* the *canonical effect* of the mutation and *δ_i_* its *actual effect*.

For enzymes *j* ≠ *i*, we have

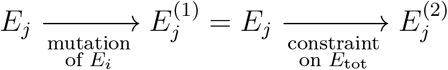

where 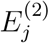 is the concentration of enzyme *j* after applying the constraint on total enzyme concentration. To find the expression of 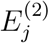, we applied the proportionality rule:

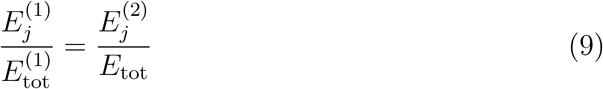

and using equation 8 we get (Supporting Information SI.B.1.3):

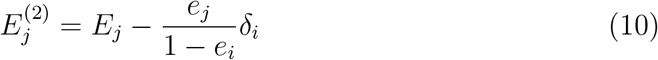

###### 3. Co-regulation and competition

If there is both competition and co-regulation between all enzymes, the two-step process is the same as above for enzyme *i* targeted by a mutation:

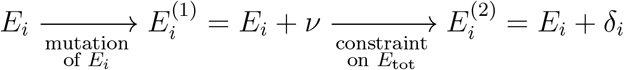

but here 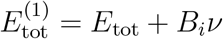 (equation 5). So applying the proportionality argument (equation 7 and Supporting Information SI.B.1.4), we get:

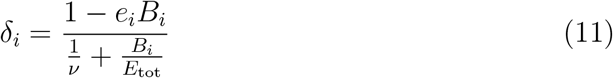

For enzymes *j* ≠ *i*, we have

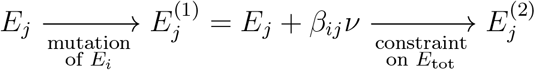

where 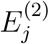 is the concentration of enzyme *j* after co-regulation *and* reduction of the total enzyme concentration from 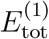 to *E*_tot_. Using equations 9 and 11, we get (Supporting Information SI.B.1.4):

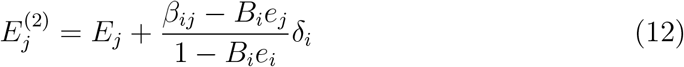

In summary, non-independence of enzyme concentrations results in changes in the concentration of enzymes that are not the primary targets of the mutation. The amplitude of concentration change of non-target enzymes depends on the cause of interdependence:

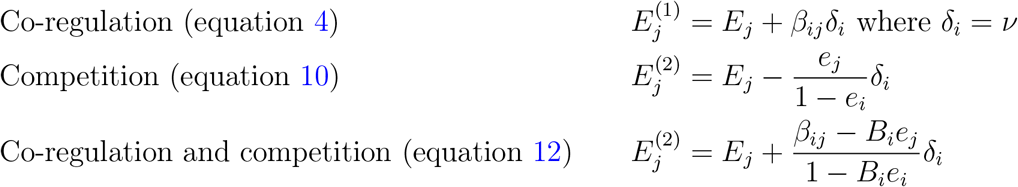

Thus the concentration of the mutant enzyme 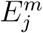 can be expressed as the sum of *E_j_* and a *pseudo-mutation* effect:

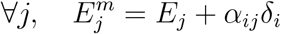

This *pseudo-mutation* effect is the product of the actual mutation effect *δ_i_* by a coefficient called the *redistribution coefficient*, noted *α_ij_* (note that ∀*i*, *α_ii_* = 1).

The general expression of the redistribution coefficient is 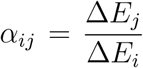, where Δ*E_j_* is the change of non-target enzyme *j* and Δ*E_i_* = *δ_i_* is the actual effect of the mutation on target enzyme *i*. Going to the limit when *δ_i_* is very small, we obtain:

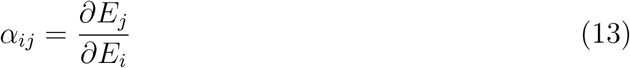

which is the redistribution coefficient *α_ij_* defined by Lion et al. (2004).

Depending on the situations, we have:

1. Independence: ∀*i, j* ≠ *i*, *α_ij_* = 0
2. Co-regulation: ∀*i, j* ≠ *i*, *α_ij_* = *β_ij_* ≠ 0
3. Competition: ∀*i, j* ≠ *i*, 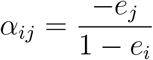
4. Co-regulation and competition: ∀*i, j* ≠ *i*, 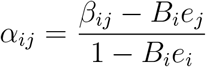

##### 2.1.1.3 Quantifying the effect of enzyme concentration variation on the flux

To quantify the effect on the flux of variation in enzyme concentration *E_i_*, we used the *combined response coefficient* (hereafter, the flux response coefficient) (de Vienne et al. 2001a)

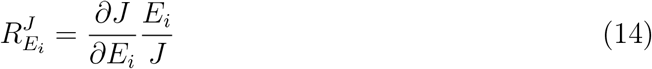

The flux response coefficient is more general than the classical flux control coefficient (Kacser et al. 1995), the latter being valid only if enzymes are independent. The flux response coefficient of enzyme *i* includes both the flux control coefficient 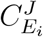 of enzyme *i* and a term that accounts for enzyme interdependence, which contains the flux control coefficients of the other enzymes and the redistribution coefficients *α_ij_* (Lion et al. 2004):

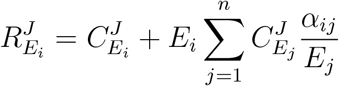

In Supporting Information SI.B.2, we present another expression of 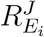 that is less intuitive but more convenient for mathematical treatment.

#### 2.1.2 The evolution model

##### 2.1.2.1 Adaptive dynamics

To analyze the evolution of enzyme concentrations, we used an adaptive dynamics approach (Brännström et al. 2013), assuming a constant environment.

We assumed that evolution proceeds by the successive fixation of mutations modifying enzyme concentration. We applied the mutual exclusion principle, *i.e.* two phenotypes cannot coexist because the mutations are assumed to be instantaneously fixed or eliminated (Dieckmann and Law 1996). On an evolutionary timescale, if a mutation produces a novel phenotype, either it is fixed, becoming the new resident, or it is eliminated. The vector of enzyme concentrations of the resident is noted 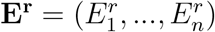 and that of the mutant 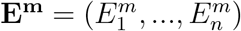.

Mutations appear at rate *μ* per time unit. They are independent and each mutation targets only one enzyme. The actual effect of a mutation on the target enzyme *i* is *δ_i_*. Target enzyme concentration after a mutation is:

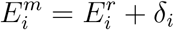

The concentration of each of the other enzymes is:

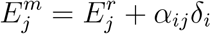

The selection coefficient of a mutation targeting enzyme *i* is:

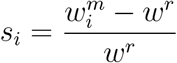

where *w* is fitness. We assumed that fitness *w* is proportional to flux *J*, so we have:

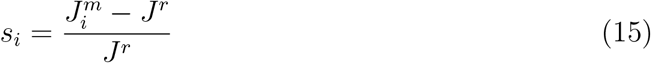

The phenotype under selection, namely the flux, is calculated following equation 1.

Since we considered a haploid population of constant effective size *N* with non-overlapping generations, the fixation probability of a mutation is (Crow and Kimura 1970):

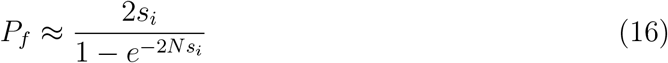

when *s_i_* is small. This expression was used in our computer simulations.

In our analytical developments, we neglected deleterious mutations because their negative selection coefficients result in negligible fixation probability. Since we assumed large population sizes (for all *i*, *Ns_i_* ≫ 1), we could use the approximation:

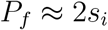

The higher the mutant flux *J^m^* is relative to the resident flux *J^r^*, the higher the fixation probability of the mutation. After fixation or loss, a new mutation appears, and the process is repeated.

##### 2.1.2.2 The differential equation system of enzyme evolution

On a continuous timescale, the variation in concentration of enzyme *i* can be represented as a stochastic process. At each time *t*, this variation is a random variable that can be described with a complete event system:

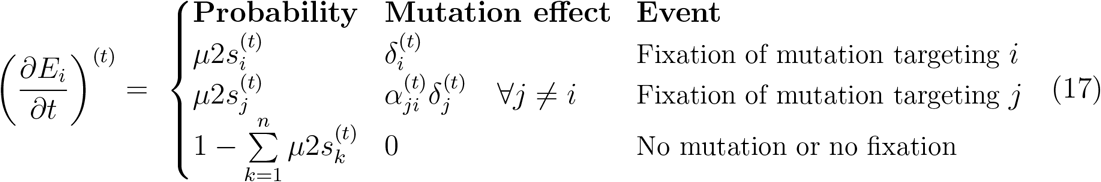

where *μ* is the probability of occurrence of a mutation, assumed to be independent of the target enzyme. Because this relation is valid at each time *t*, we will not use the superscript *t* thereafter. At each time, variation in *E_i_* can be caused by the fixation of either a mutation affecting enzyme *i* (probability 2*s_i_*) or a mutation affecting any other enzyme (probability 2*s_j_*), with mutation effect *δ_i_* and *α_ji_δ_j_*, respectively. If none of these events happen (probability 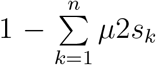), *E_i_* does not change. Therefore, the *average* variation in concentration of enzyme *i* at each time *t* is:

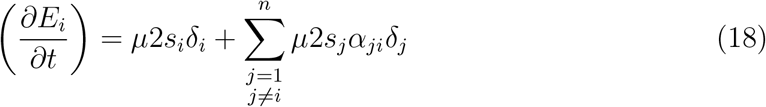

This relation is valid at all times *t*, and because *α_ii_* = 1, we can write:

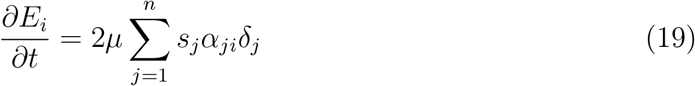

This differential system allowed us to search for steady states – called evolutionary equilibria – of enzyme concentrations when there is selection for increased flux. When enzymes are independent, *∂E_i_/∂t* s strictly positive, and thus enzyme concentrations increase indefinitely and do not reach equilibrium. Consequently, we searched for evolutionary equilibria of *relative* enzyme concentrations, which is given by:

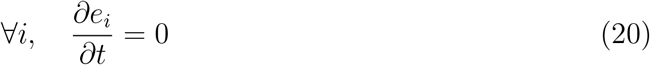

##### 2.1.2.3 Selective neutrality of enzyme concentration variation

We addressed the question of the neutrality of the variations in enzyme concentrations near evolutionary equilibrium. The “neutral zone” is the area where the absolute value of the selection coefficient is close to zero – classically under a threshold of 1*/N* for haploid populations (Kimura 1985).

To analytically study the way constraint-dependent factors influence the neutral zone at evolutionary equilibrium, we used an expression of *s_i_* that explicitly contains the three relevant variables, *i.e.* 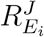, *δ_i_* and *E_i_*. Assuming that the mutation effect *δ_i_* is small compared to the concentration *E_i_* (*δ_i_* ≪ *E_i_*), we can write 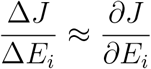. Therefore, from equation 14, the selection coefficient is given by (Supporting Information SI.B.3.1):

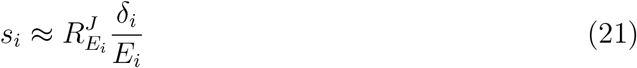

The size of the neutral zone determines what we call the *range of neutral variation* (RNV) of the concentration, the amplitude of which is:

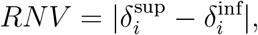

where 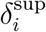 (resp. 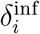) is the actual effect of a mutation such that 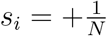 (resp. 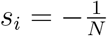), *s_i_* being the superior (resp. inferior) limit of the neutral zone. To determine the *δ_i_*’s, we used the initial equation 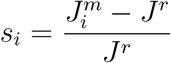 (equation 15), where *J^r^* is the resident phenotype and 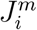 the mutant phenotype following a mutation targeting enzyme *i*. From equation 1, and after transformations, we obtain (Supporting Information SI.B.3.2):

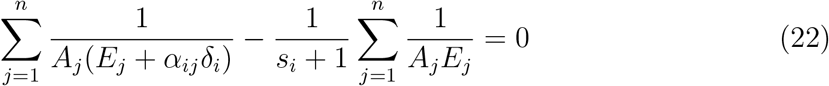

The solutions of this equation for *s_i_* = +1*/N* and *s_i_* = − 1*/N* give 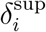 and 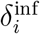, respectively. Due to the complexity of the resulting equations, we wrote a script that numerically computes the limits 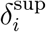 and 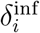 of the RNVs for different scenarios of enzyme interdependence (function `uniroot of the R Stats` package [(R Development Core Team 2008)]).

### 2.2 Computer simulations

To assess the validity of the mathematical predictions, we performed Markov Chain Monte-Carlo simulations using the R software (R Development Core Team 2008). We considered haploid populations of size *N* = 1 000 evolving in a constant environment (*X* = 1). At a given time step, all individuals in the population are identical, *i.e.* have the same phenotype *J* and the same enzyme concentrations (the “genotype”) **E** = (*E*_1_, *E*_2_, …*E_n_*) and the same pseudo-activities **A**= (*A*_1_, *A*_2_, …*A_n_*). Simulations were performed for a three-enzyme metabolic pathway (*n* = 3), with invariable pseudo-activities *A*_1_ = 1, *A*_2_ = 10 and *A*_3_ = 30 and an initial total enzyme concentration 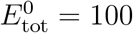 in arbitrary units (see Figure 1A).

**Figure 1.**
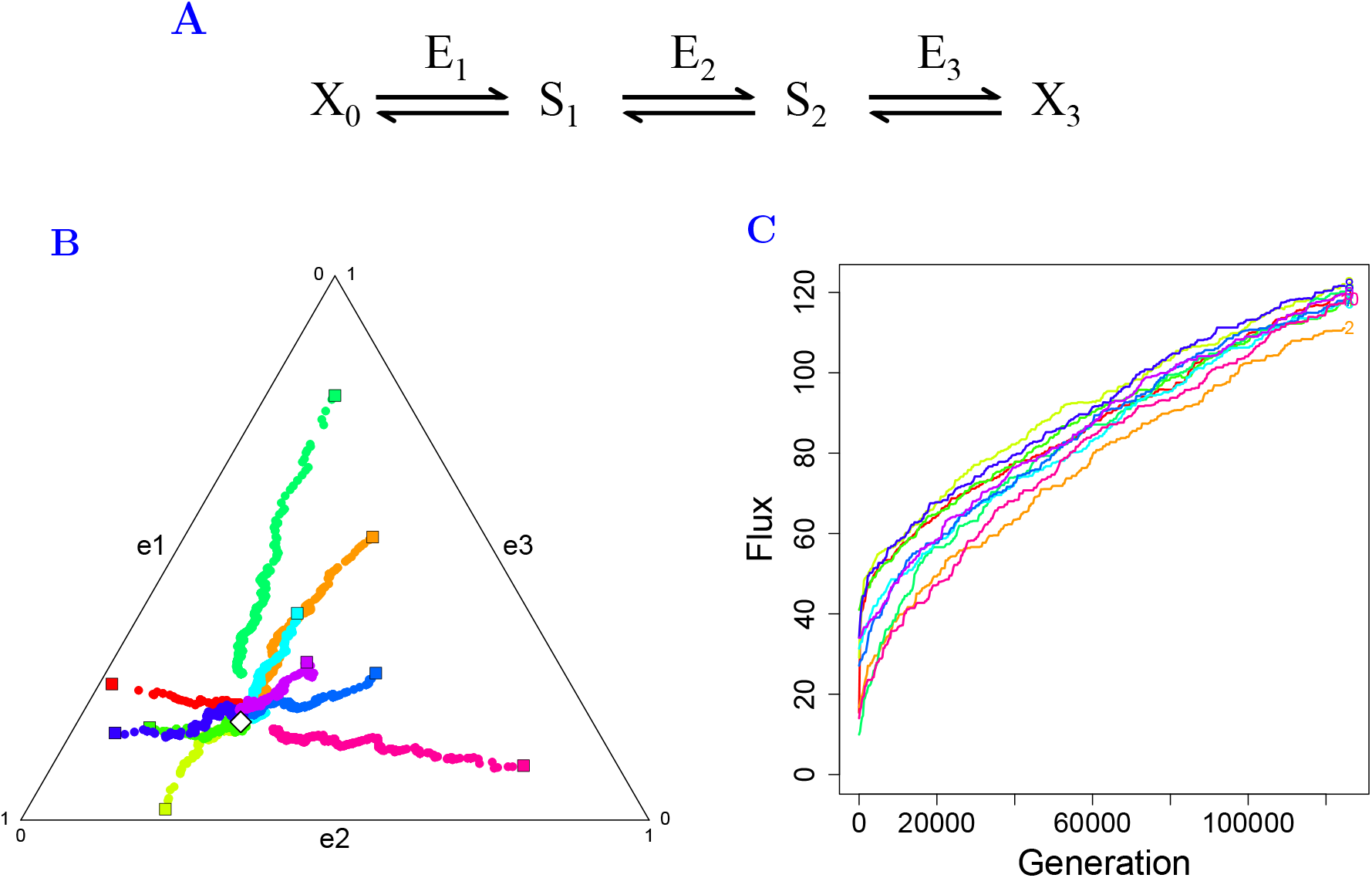
Simulation of enzyme evolution in a three-enzyme pathway when enzyme concentrations vary independently. (A) The three-enzyme pathway used in the simulations. Parameters values are: *X* = 1, *A*_1_ = 1, *A*_2_ = 10, *A*_3_ = 30, 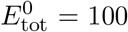, *N* = 1 000. (B) Evolutionary trajectories of relative enzyme concentrations *e*_1_, *e*_2_ and *e*_3_ represented in a triangular diagram for ten simulations (one color per simulation) with different initial enzyme concentrations (colored squares). The relative enzyme concentrations converge towards the theoretical equilibrium **e**\* (white diamond). (C) Evolution of the flux over the course of generations from simulations shown in B.

The vector of initial concentrations **E**^0^ was changed for each simulation. Initial concentrations 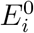 were drawn in 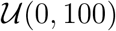, and redistributed proportionally to have 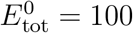. *E*_tot_ varied over time in the case of independence and co-regulation, but was fixed at *E*_tot_ = 100 in the case of competition.

Co-regulation coefficient values were *β*_12_ = 0.1, *β*_23_ = 2 and hence *β*_13_ = 0.2 for the positive co-regulations and *β*_12_ = 0.32, *β*_23_ = −0.43 and hence *β*_13_ ≈ −0.138 for the negative co-regulations. The resulting co-regulation matrices **M**_*β*_ were computed (Supporting Information SI.B.1.1).

At each time step, a mutation randomly affected one of the enzymes. The mutation effect *ν* was drawn in 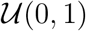; the sign of *ν* was randomly drawn (1:1 ratio). Thus the concentration of target enzyme *i* changed from 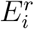 to 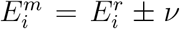; the concentration of all enzymes varied according to the type of dependence considered (see the two-steps process described in paragraph 2.1.1.2). If a concentration became negative, it was set to zero. Finally, we computed the values of the flux, the selection coefficient and the fixation probability of the mutant (equations 1, 15 and 16, respectively).

To determine whether a mutation was fixed or lost, we drew a number in 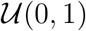, and if this number was smaller than *P_f_* the mutation was fixed, otherwise it was lost. If the mutation was fixed, the mutant became the new resident, *i.e.* all individuals displayed the same genotype and phenotype. If the mutation was lost, genotype and phenotype were unchanged. We then proceeded to the next time step. Thus, a time step is the interval between two successive mutations. Populations evolved over 125,000 time steps, and the system values were recorded every 250 steps.

These simulations allowed us to relax some of the assumptions of the mathematical analysis, namely small mutation effects, large population size and the absence of fixation of deleterious mutations.

#### Data and code availability

Scripts of the simulations and custom functions are written in the R language (R Development Core Team 2008). Along with the simulation data, they are included in a home-made package called `SimEvolEnzCons`.

## 3 Results

We have developed a theoretical framework that describes the evolution of enzyme concentrations in a pathway under selection for increased metabolic flux. We considered that: (i) the flux can be modified by mutations targeting any enzyme in the pathway; (ii) variation in enzyme concentrations can be not independent as a result of competition for cellular resources and/or co-regulation. We based our developments on the principles of adaptive dynamics (Brännström et al. 2013), using the enzyme-flux relationship of the Metabolic Control Theory (MCT) (Kacser and Burns 1973; Heinrich and Rapoport 1974; Kacser et al. 1995) expanded by Heinrich et al. (1991) and Lion et al. (2004) to take into account possible dependencies between enzyme concentrations.

Flux *J* through a linear metabolic pathway of *n* Michaelian enzymes was expressed as a function of the parameters of all enzymes (equation 1 in Material & Methods):

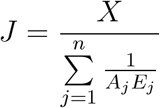

where *A_j_* is the *pseudo-activity* of enzyme *j*, assumed to be constant, and *E_j_* is the concentration of enzyme *j* that can be affected by mutation (see Glossary of mathematical symbols in appendix A).

If there is selection for increased flux, the evolutionary rate of enzyme concentrations can be expressed by a differential equation system (equation 19), allowing us to: (i) search for steady states of relative enzyme concentrations *e_i_* (∀_*i*_, *∂e_i_/∂t* = 0), which we call *evolutionary equilibria*; (ii) analyze flux variation during the evolutionary process.

To assess the validity of our analytical results, we performed simulations of long-term evolution using less stringent assumptions than in the analytical approach.

Finally, we analyzed the extent to which selection for increased metabolic flux results in the selective neutrality of mutations targeting enzyme concentrations.

### 3.1 Equilibria of relative enzyme concentrations under directional selection for increased flux

We searched for evolutionary equilibria of relative enzyme concentrations in four situations: independence between enzyme concentrations, fixed total enzyme concentration (due to competition), co-regulation between all enzyme concentrations and both competition and co-regulation.

#### 3.1.1 Case 1: Independence between enzymes

When the enzyme concentrations are independent, the evolutionary equilibrium of relative enzyme concentrations is (Supporting Information SI.B.4):

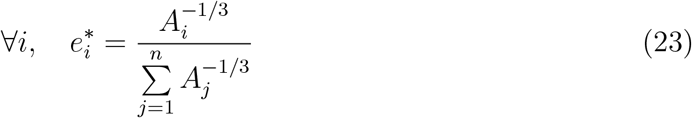

Simulation results of a three-enzyme pathway (Figure 1A) are consistent with this theoretical equilibrium: the evolutionary trajectories converge towards the point of the space of relative concentrations defined by equation 23 (Figure 1B). At this point, the flux response coefficients, which quantify the sensitivity of the flux to variation in enzyme concentrations, are (Supporting Information SI.B.4):

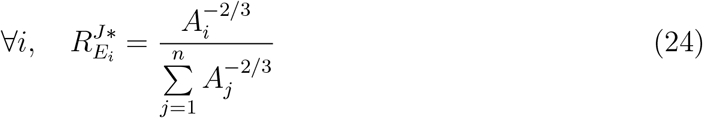

Because the flux response coefficients are strictly positive and there is no upper limit on enzyme concentration, the flux is continuously increasing (Figure 1C) while the proportions of the different enzymes remain constant at equilibrium (equation 23).

From equations 23 and 24, we see that the relative concentrations and the flux response coefficients at equilibrium are inversely related to the pseudo-activities: enzymes with low pseudo-activity have a high concentration relative to others enzymes and exert a high control on the flux. For these enzymes, variation in abundance is less likely to be selectively neutral (see Section 3.2.2).

#### 3.1.2 Case 2: Competition for cellular resources

In order to model the effects of limited cellular resources, the total enzyme abundance *E*_tot_ in the pathway was considered to be constant. When a mutation targets the concentration of an enzyme, the concentrations of all enzymes are assumed to be redistributed proportionally to maintain constant the total concentration. As a consequence, the relationship between relative enzyme concentrations and flux is a dome in the multidimensional space of relative enzyme concentrations, the shape of which is determined by the vector of pseudo-activities **A** (Figure 2A). The flux cannot exceed a maximum value *J*_max_, which is the flux value at evolutionary equilibrium.

**Figure 2.**
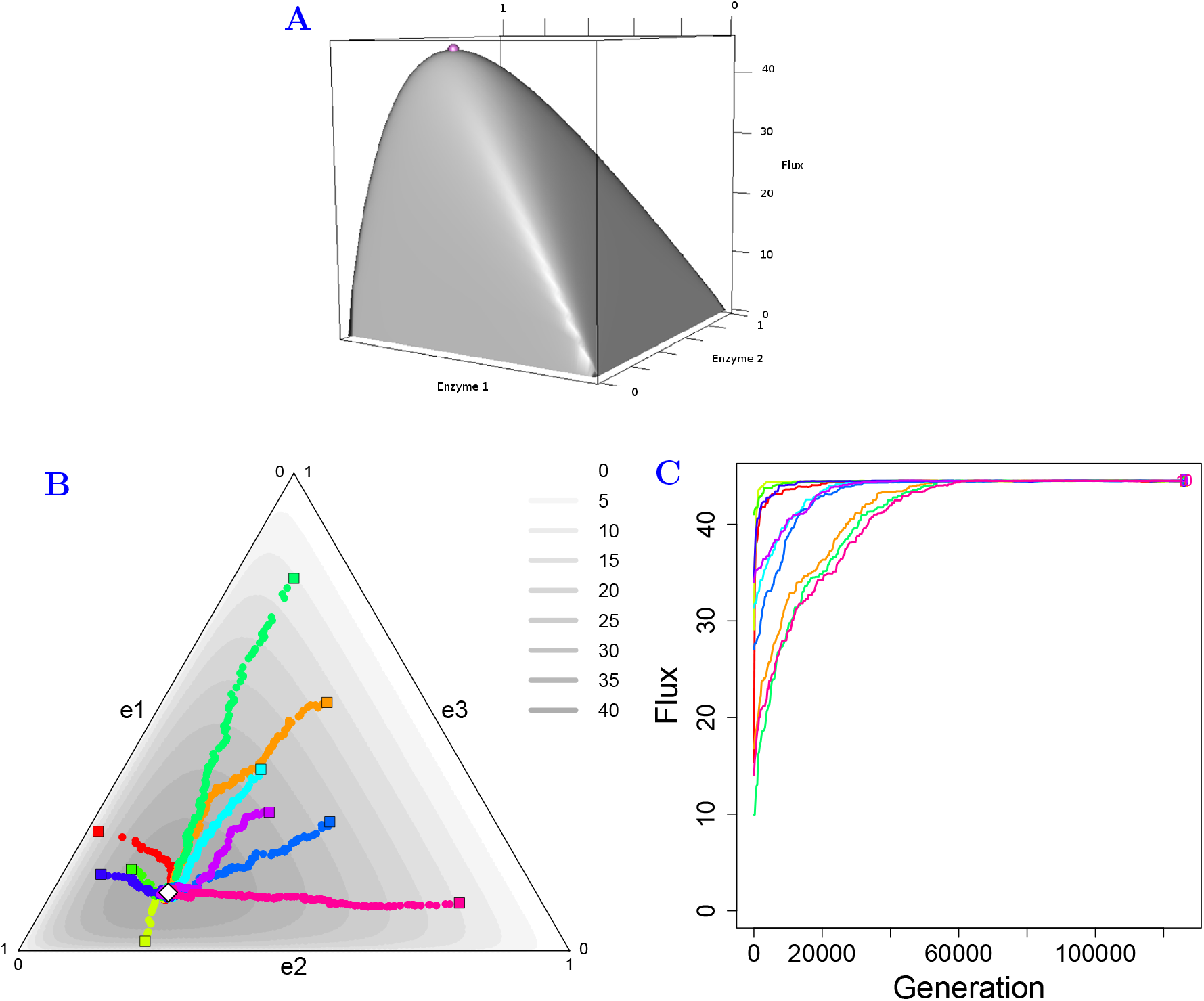
Simulation of enzyme evolution in a three-enzyme pathway when there is competition. (A) The flux-enzyme relationship is dome-shaped because *E*_tot_ is constant. The purple point is the maximal flux *J*_max_, corresponding to the evolutionary equilibrium. (B) and (C) Representations and symbols are as in Figure 1; the grayscale in (B) corresponds to the flux values. When there is competition, the relative enzyme concentrations converge towards the equilibrium (white diamond) and the flux is limited. Parameter values are: *X* = 1, *A*_1_ = 1, *A*_2_ = 10, *A*_3_ = 30, 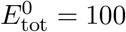, *N* = 1 000.

At *J*_max_ the flux response coefficients are:

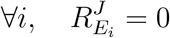

because at the top of the dome, the derivative of the flux with respect to *E_i_* is null in all dimensions. From this equality, the evolutionary equilibrium of relative enzyme concentrations at *J*_max_ can be found (Supporting Information SI.B.5):

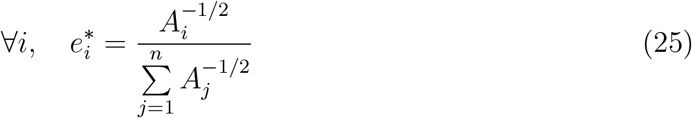

As in the case of independence, simulation results were consistent with this theoretical equilibrium (Figure 2B). As previously, the higher the pseudo-activity of the enzyme, the lower its relative concentration, but due to the difference in the value of the power of *A_i_* (−1/2 instead of −1/3), the relative enzyme concentrations at evolutionary equilibrium were more dispersed when there was competition than when there was independence.

From equation 25, we can calculate *J*_max_, which is linearly related to *E*_tot_ (Supporting Information SI.B.5):

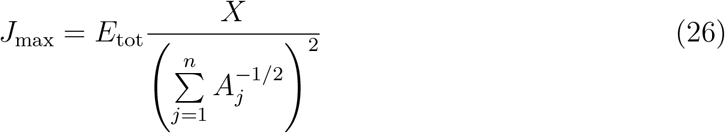

Figure 2C shows the results of the simulations of flux evolution when *E*_tot_ is fixed. The expected *J*_max_ was reached whatever the value of the initial enzyme concentrations.

#### 3.1.3 Case 3: Co-regulation between enzyme concentrations

When there is co-regulation between enzyme concentrations, in the absence of competition, a mutation targeting a given enzyme results in variation in the concentration of the other enzymes. To model this phenomenon, we used the *co-regulation coefficient β_ij_* – the slope of the linear relation between *E_i_* and *E_j_* –, which quantifies the variation in *E_j_* when a mutation affects *E_i_*.

When all enzyme concentrations are related to each other (∀(*i, j*), *β_ij_* ≠ 0), the evolutionary equilibrium of the relative enzyme concentrations is (Supporting Information SI.B.6.1.1):

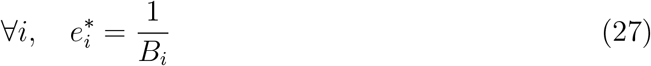

where 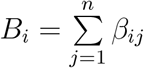, the *global regulation coefficient* of enzyme *i*, quantifies the effect of the variation of *E_i_* on the total enzyme concentration (equation 6): *B_i_* = Δ*E*_tot_/Δ*E_i_*. This equilibrium has a non-intuitive property: at this point, mutations, whatever their effect, no longer modify the *relative* enzyme concentrations (Supporting Information SI.B.6.1.2). Metaphorically, we will call this point an *evolutionary black hole*.

The enzyme concentrations are assumed to be linearly related. Consequently, in the multidimensional space of relative enzyme concentrations, the system can be described as a point moving on a straight line *ε*. The parametric equation of *ε* can be determined from two known points, the initial relative enzyme concentrations **e^0^** and the theoretical equilibrium **e**\* (Supporting Information SI.B.6.2):

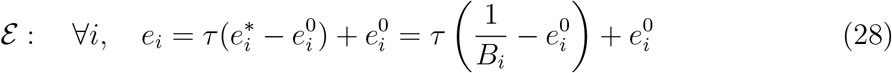

where *τ* is the driving variable. When *τ* = 0, the system is at the initial state **e^0^** and when *τ* = 1, the system is at the theoretical equilibrium **e**\*. For a given **e^0^** and **M**_*β*_ (the matrix of all pairwise co-regulation coefficients), the relative concentrations **e** are strictly determined by the value of *τ*. Consistently, results of the simulations showed linear evolutionary trajectories converging towards the theoretical equilibrium (Figures 3A and 3C), with a clear difference between positive and negative co-regulation:

##### 1. Positive co-regulation

When all *β_ij_*’s are strictly positive, the relative enzyme concentrations converge towards the theoretical equilibrium **e**\* located within the space of relative enzyme concentrations (Figure 3A). At this point, the flux response coefficients are (Supporting Information SI.B.6.1.3):

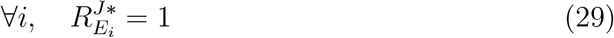

meaning that every enzyme strictly controls the flux. This is not in contradiction with the previously mentioned evolutionary black hole, because the flux response coefficients concern the *absolute* enzyme concentrations, which increase indefinitely, as does the flux (Figure 3B), while the *relative* enzyme concentrations are stuck at the theoretical equilibrium.

**Figure 3.**
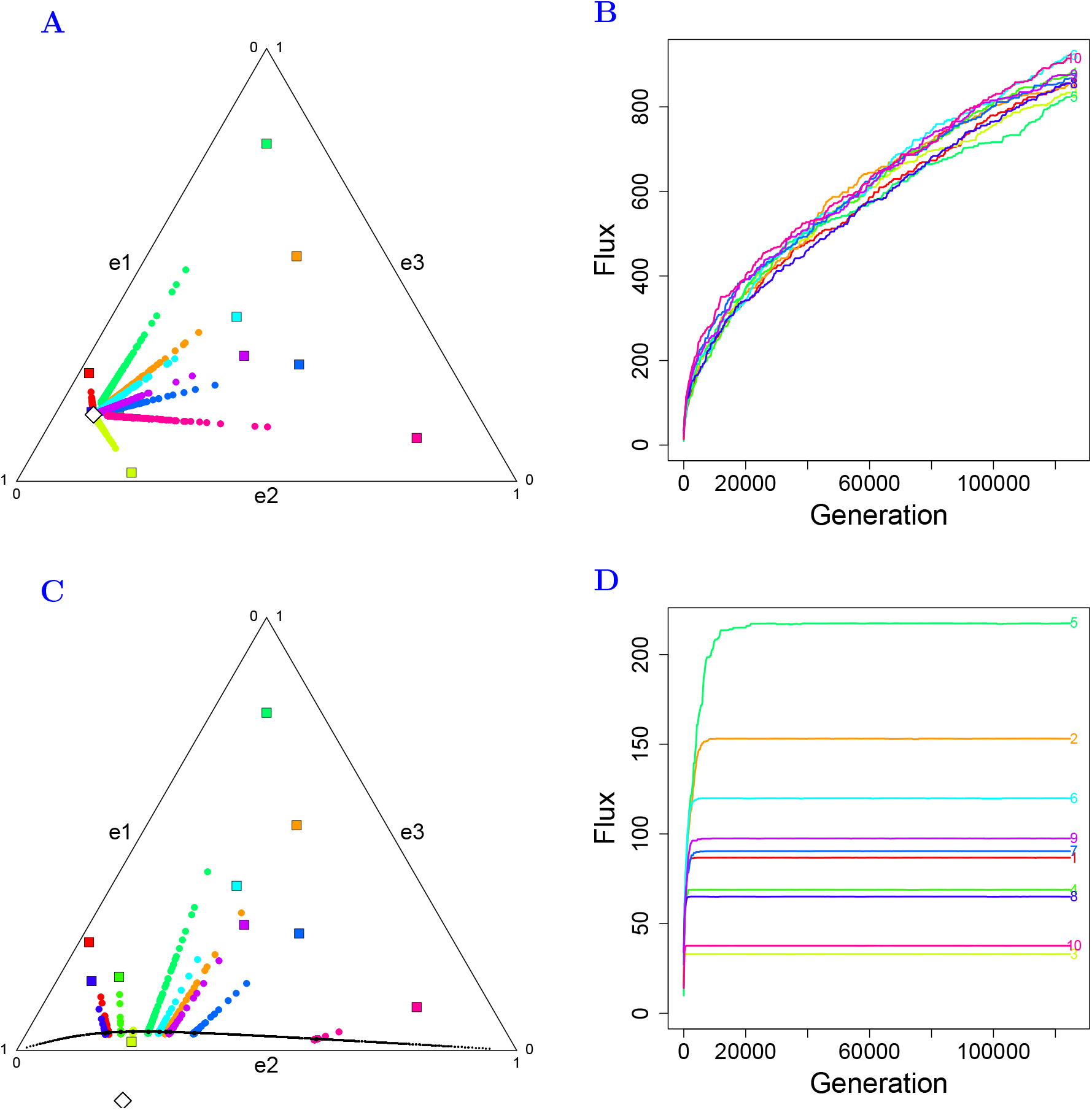
Simulation of enzyme evolution in a three-enzyme pathway when there is co-regulation. Representations and symbols are as in Figure 1. (A) and (B) Positive co-regulation. (C) and (D) Negative co-regulation. When there is co-regulation, the evolutionary trajectories in (A) and (C) are linear. In the case of negative co-regulation, the theoretical equilibrium **e**\* (white diamond in (C)) is outside the triangle. The black curve in (C) represents the sets of points of the effective equilibrium 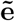 determined numerically for a large number of initial concentrations **e^0^**. The relative enzyme concentrations fluctuate on both sides of this curve due to the mutation-selection-drift balance. (B) and (D) Evolution of the flux over the course of generations from simulations shown in (A) and (C), respectively. Parameter values are: *X* = 1, *A*_1_ = 1, *A*_2_ = 10, *A*_3_ = 30, 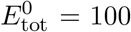, *N* = 1 000, *β*_12_ = 0.1, *β*_23_ = 2 (A and B) or *β*_12_ = 0.32, *β*_23_ = *−*0.43 (C and D).

##### 2. Negative co-regulation

If at least one co-regulation coefficient is negative, then at least one enzyme has a negative global regulation coefficient *B_i_* (Supporting Information SI.B.1.2). As concentrations cannot be negative, the theoretical equilibrium **e**\* is unattainable because it is outside the space of possible relative enzyme concentrations (Figure 3C). There is another evolutionary equilibrium, which we called the *effective equilibrium* 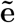, defined by (Supporting Information SI.B.6.3):

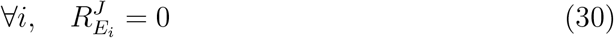

Null response coefficients mean that in this case there is a maximum value of the flux, 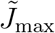, for the same reason as in the case of competition: the negative co-regulation coefficients prevent the flux from exceeding a certain value because mutations that increase (resp. decrease) the concentration of an enzyme will decrease (resp. increase) the concentration of the other enzyme(s).

The *effective equilibrium* 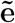 could not be found analytically but could be found numerically from another expression of equation 30 (Supporting Information SI.B.6.3):

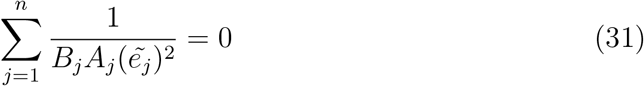

The black curve on Figure 3C shows the set of solutions 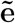 satisfying equation 31 for a given set of *A_i_* and *B_i_* values and a large number of initial enzyme concentrations **e^0^**. All evolutionary trajectories converge towards the theoretical equilibrium, but stop on this curve at their respective effective equilibrium. Figure 3D shows the evolution of the corresponding fluxes and their respective 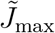.

#### 3.1.4 Case 4: Competition *plus* co-regulation

If there is competition and co-regulation, the relationship between flux and enzyme concentrations is a dome in the multidimensional space of relative enzyme concentrations, but here the relative enzyme concentrations are constrained by the effects of co-regulation to follow line *ε* (equation 28). Consequently, the flux evolves following the curve defined by the intersection of the dome with the upright plane drawn from line *ε* in space (**e**, *J*) (Figure 4). There are three remarkable points on this curve: (i) the flux value corresponding to the initial relative enzyme concentrations **e^0^**(black point); (ii) the flux value corresponding to the theoretical equilibrium of relative enzyme concentrations **e**\* (red point); (iii)the maximum of this curve, 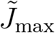, corresponding to the effective equilibrium of relative enzyme concentrations 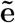 (yellow point). Note that for a given **e**\*, the orientation of the plane is determined by **e^0^**.

**Figure 4.**
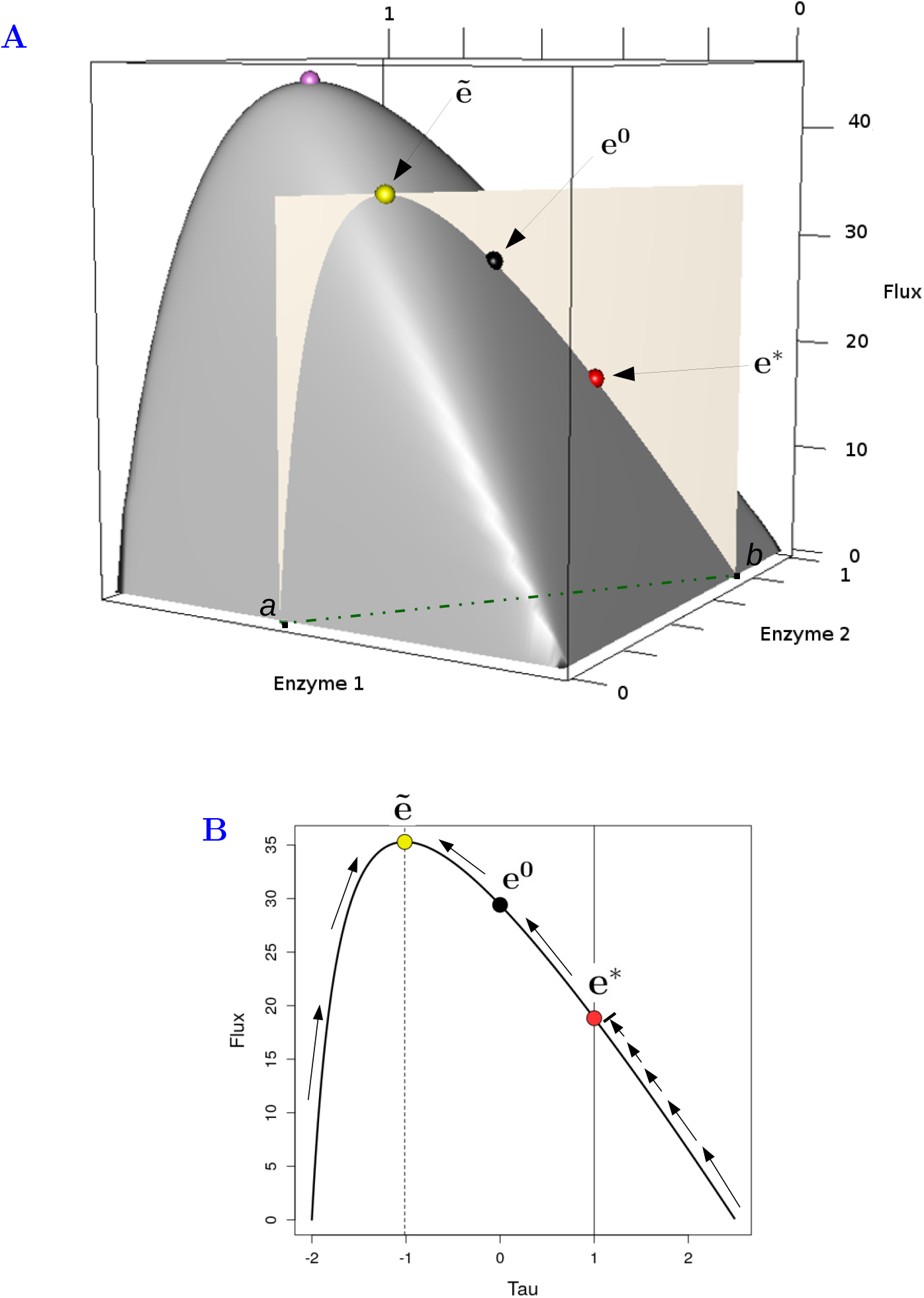
Relationship between flux and relative enzyme concentrations when there is both competition and positive co-regulation in a three-enzyme pathway. (A) When there is only competition, the relation between flux and relative concentrations of enzymes 1 and 2 is represented by a dome. The purple point indicates the maximum flux *J*_max_. The ivory section represents the upright plane drawn from line along which enzyme concentrations vary when there is positive co-regulation (green dashed line on the plane). Red point: flux value at theoretical equilibrium **e**\*. Black point: flux value at the initial enzyme concentrations **e^0^**. Yellow point: maximum flux value 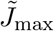 at effective equilibrium 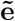. (B) Relationship between *J* and *τ*, the driving variable of line *ε*, corresponding to the ivory section of the dome in (A). Solid vertical line: abscissa of the theoretical equilibrium. Dashed vertical line: abscissa of the effective equilibrium corresponding to 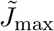. Colored points: same as in (A). The arrows indicate the variation of flux under selection for different initial enzyme concentrations. Note that the red point (the theoretical equilibrium) is a singular point that cannot be passed if the initial flux is lower than the flux at **e**\* *and* if **e**\* and **e^0^** are on the same side of the curve. Parameter values are: *X* = 1, *A*_1_ = 1, *A*_2_ = 10, *A*_3_ = 30, 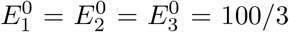, *β*_12_ = 2.5, *β*_23_ = 0.6, corresponding to *B*_1_ = 5, *B*_2_ = 2 and *B*_3_ = 3.33.

The theoretical equilibrium is the same as in the case of co-regulation alone (Supporting Information SI.B.7.1.1):

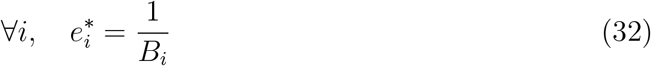

and the effective equilibrium is also defined by (Supporting Information SI.B.7.2):

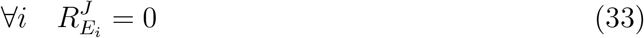

which leads to the following equation:

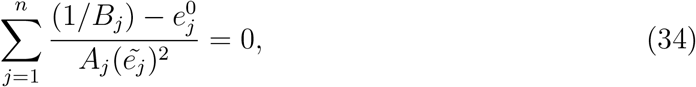

from which 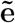 can be found numerically. If we consider a large number of possible **e^0^**’s for given **M**_*β*_ and **A**, we can compute a set of points corresponding to 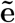, which defines a closed curve in the case of positive co-regulation (Figure 5A) and an open curve in the case of negative co-regulation (Figure 5C). As previously, all evolutionary trajectories end on these curves. At 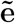 the flux reaches 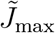, which depends on **e^0^**, **M**_*β*_, **A** and 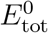 (Figures 5B and 5D).

**Figure 5.**
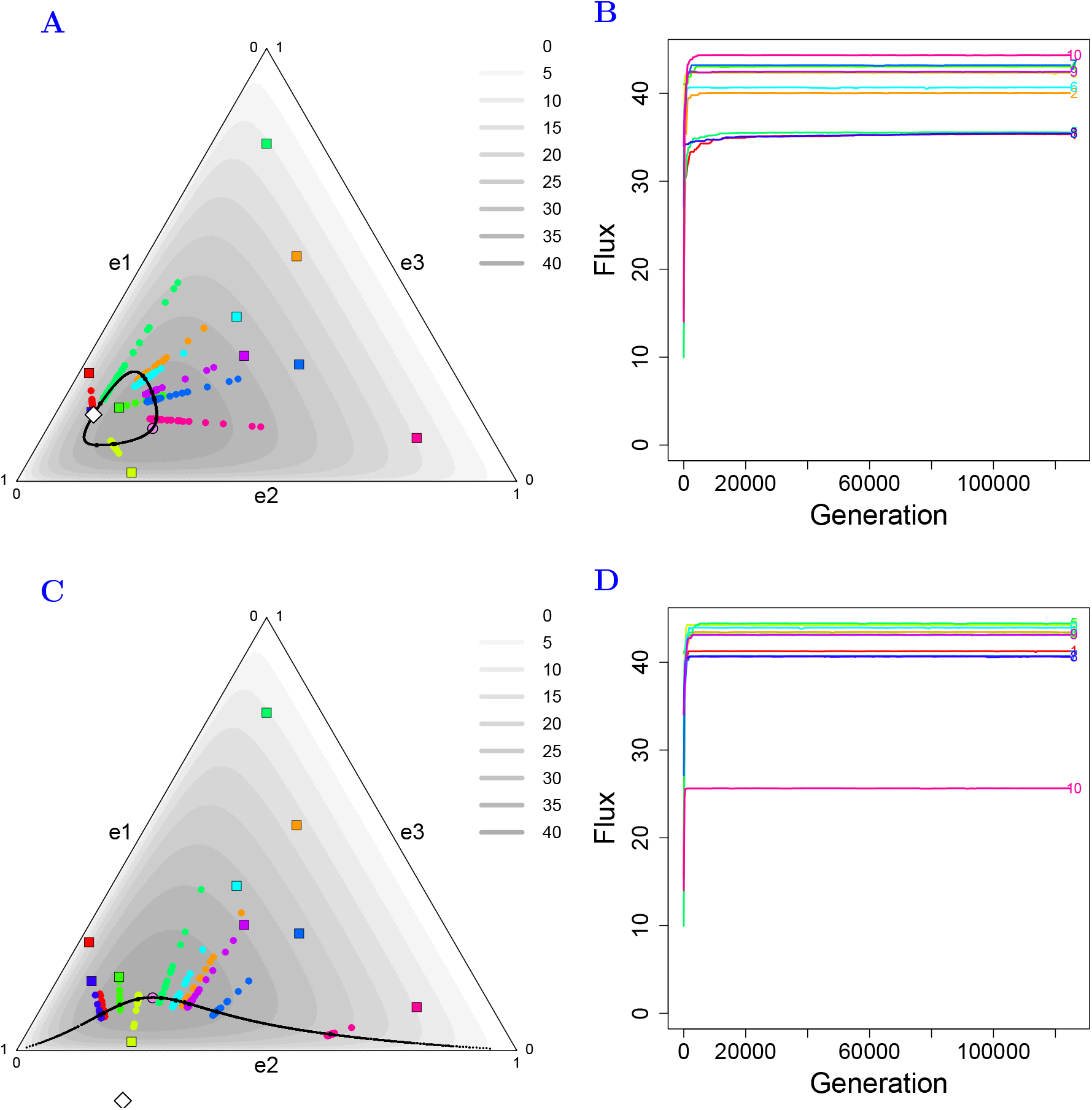
Simulation of enzyme evolution in a three-enzyme pathway when there is competition and co-regulation. Representations and symbols are as in Figure 1. (A) and (B) Competition plus positive co-regulation. (C) and (D) Competition plus negative co-regulation. In (A) and (C), the grayscales correspond to the flux values and the black curves represent the sets of points of the effective equilibrium determined numerically for a large number of initial concentrations **e^0^**. The evolutionary trajectories converge towards, but do not reach, the theoretical equilibrium *e*\* (white diamonds). They fluctuate on both sides of the curves due to the mutation-selection-drift balance. (B) and (D) The evolution of the flux over the course of generations from the ten simulations shown in A and C, respectively. Parameter values are: *X* = 1, *A*_1_ = 10, *A*_2_ = 10, *A*_3_ = 30, 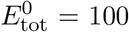, *N* = 1 000, *β*_12_ = 0.1, *β*_23_ = 2 (A and B) or *β*_12_ = 0.32, *β*_23_ = −0.43 (C and D).

When there is both competition and positive co-regulation, two possible evolutionary equilibria exist, depending on the initial relative enzyme concentrations **e^0^** (Figure 4). If the theoretical equilibrium is reached first, the evolutionary black hole, which is also valid in this situation (Supporting Information SI.B.7.1.2), will prevent any flux increase despite the selective pressure. If the effective equilibrium is reached first, selection for high flux will prevent the theoretical equilibrium from being reached because the flux value is higher at the effective equilibrium than at the theoretical equilibrium.

#### 3.1.5 Summary of the results on evolutionary equilibria

In summary, the conclusions drawn from the analytical developments and the simulations are as follows:

- Independence: The flux increases continuously but the relative enzyme concentrations reach an equilibrium value that depends on the pseudo-activities of the enzymes.
- Competition: The flux increases up to a maximum value that depends on both the pseudo-activities of the enzymes and the total enzyme concentration. The relative enzyme concentrations reach an equilibrium that depends on the pseudo-activities of the enzymes, but which differs from the one obtained in the case of independence.
- Co-regulations without competition: The relative enzyme concentrations tend towards an equilibrium that depends only on the co-regulation coefficients between enzymes. If the co-regulation coefficients are all positive, the flux increases continuously and equilibrium is reached. This equilibrium is an evolutionary black hole because, once it is reached, additional mutations cannot change the *relative* enzyme concentrations. If there is at least one negative co-regulation coefficient, the flux evolves towards a maximum and a different equilibrium is reached, called the effective equilibrium, which depends not only on the co-regulation coefficients and the pseudo-activities of the enzymes, but also on the initial enzyme concentrations.
- Competition *plus* co-regulation: This case is qualitatively similar to the case with negative co-regulation alone, with a flux maximum and a multi-factor effective equilibrium. The difference resides in the shape of the adaptive landscape.

Tables 1 and 2 summarize the evolutionary equilibria and the limit to the flux value in the different situations.

**Table 1.** Theoretical *e*\* and effective 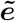 equilibrium for relative enzyme concentrations.

|  | Equilibrium | Without competition | With competition |
| --- | --- | --- | --- |
| **Without co-regulation** | Theoretical | $\forall i, e_i^* = \frac{A_i^{-1/3}}{\sum_{j=1}^n A_j^{-1/3}}$ | $\forall i, e_i^* = \frac{A_i^{-1/2}}{\sum_{j=1}^n A_j^{-1/2}}$ |
| **Positive co-regulation** | Theoretical | $\forall i, e_i^* = 1/B_i$ | $\forall i, e_i^* = 1/B_i$ |
| | Effective | — | $\tilde{e} = f(\mathbf{e}^0, \mathbf{M}_\beta, \mathbf{A})$ |
| **Negative co-regulation** | Theoretical | $\forall i, e_i^* = 1/B_i$<br>cannot be reached | $\forall i, e_i^* = 1/B_i$<br>cannot be reached |
| | Effective | $\tilde{e} = f(\mathbf{e}^0, \mathbf{M}_\beta, \mathbf{A})$ | $\tilde{e} = f(\mathbf{e}^0, \mathbf{M}_\beta, \mathbf{A})$ |

**Table 2.** Limit to the flux value. The maximum value of the flux corresponds to the effective equilibrium.

|  | Without competition | With competition |
| --- | --- | --- |
| **Without co-regulation** | $J \rightarrow \infty$ | $J_{\max} = f(\mathbf{A}, E_{\text{tot}}^0)$ |
| **Positive co-regulation** | $J \rightarrow \infty$ | $\tilde{J}_{\max} = f(\mathbf{e}^0, \mathbf{M}_\beta, \mathbf{A}, E_{\text{tot}}^0)$ |
| **Negative co-regulation** | $\tilde{J}_{\max} = f(\mathbf{e}^0, \mathbf{M}_\beta, \mathbf{A}, E_{\text{tot}}^0)$ | $\tilde{J}_{\max} = f(\mathbf{e}^0, \mathbf{M}_\beta, \mathbf{A}, E_{\text{tot}}^0)$ |

### 3.2 The range of selective neutrality of enzyme concentrations

#### 3.2.1 Selective neutrality at evolutionary equilibrium

In addition to searching for evolutionary equilibria, another key question from an evolutionary point of view is the extent to which mutations affecting enzyme concentrations become neutral under selection, *i.e.* have a negligible effect on the flux, and by extension on fitness. The classical criterion defining the neutral zone for haploid populations is a value of the selection coefficient *s_i_* between −1/*N* and +1/*N* (Kimura 1985).

In our model, the selection coefficient is related to *δ_i_*, 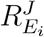 and *E_i_* following the relation 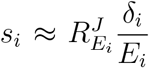 (equation 21). When *J* is limited (when there is competition, negative co-regulation or both competition and co-regulation), 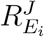 is null at the effective equilibrium of relative enzyme concentrations, and thus mutations are nearly neutral near this point. When *J* is not limited (when enzymes are independent or when there is positive co-regulation), response coefficients are constant at the theoretical equilibrium and 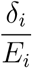 tends towards zero because *E_i_*’s increase indefinitely, and thus mutations are nearly neutral at this point. Therefore, selection for high flux leads to the near neutrality of mutations affecting any enzyme in the pathway in all situations considered.

#### 3.2.2 The range of neutral variation depends on the enzymes and the constraints

Even though the mutations affecting enzyme concentrations become neutral as relative enzyme concentrations approach equilibrium, the size of the neutral zone of mutations is expected to vary depending on the enzyme properties and the constraints applied. The zone of selective neutrality determines the *range of neutral variation* (RNV) of the concentrations. The RNV size is given by 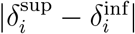, where 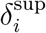 (resp. 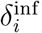) is the effect of a mutation such that 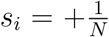 (resp. 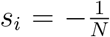), *s_i_* being the superior (resp. inferior) limit of the neutral zone. Let us examine the four situations.

##### 3.2.2.1 Case 1: Independence

From equations 21, 23 and 24, we deduced the selection coefficient at theoretical equilibrium:

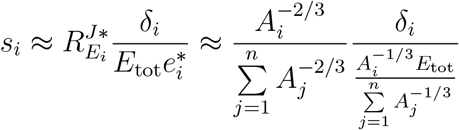

Since *δ_i_* = *ν* when enzyme concentrations are independent, it comes:

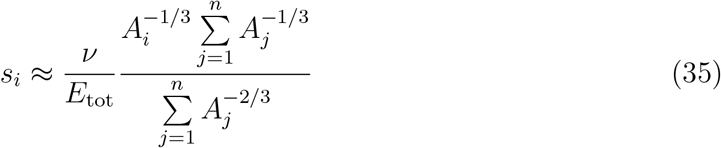

We see that, for a given ratio *ν/E*_tot_, the pseudo-activities *A_j_* alone determine the *s_i_*’s. For two enzymes *i* and *i*′, if *A_i_* < *A_i′_*, then |*s_i_*| > |*s_i′_*|, *i.e.* the selection coefficients are inversely related to the pseudo-activities. Therefore, to get out of the neutral zone, a larger mutation effect is required for enzymes with large *A_j_*’s than for those with small *A_j_*’s (Figure S1A). This means that RNV size is positively related to the pseudo-activities **A**, and hence is inversely related to the theoretical equilibria *e*\*. Numerical application was performed for a ten-enzyme pathway using equation 22 to compute the limits of the RNVs. Results were fully consistent with the theoretical predictions (Figure 6A).

**Figure 6.**
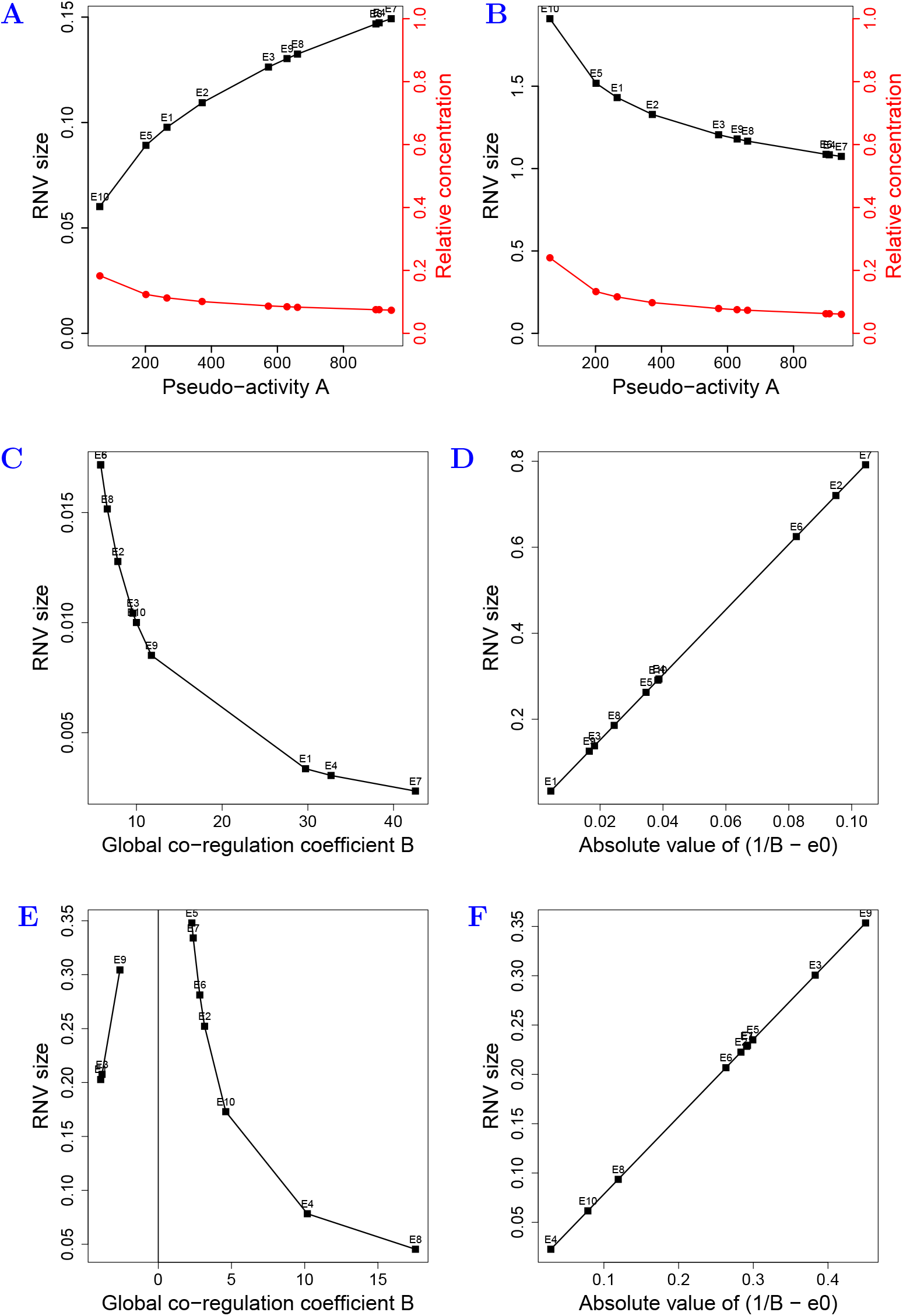
Constraint-dependent factors impacting the range of neutral variation (RNV) of enzyme concentrations. Numerical applications for a ten-enzyme pathway. If enzymes vary independently (A), the RNV size at evolutionary equilibrium is positively related to the pseudo-activities – hence negatively to the relative enzyme concentrations –, whereas the opposite is found when there is competition (B). For a given total enzyme concentration, the RNV size is one order of magnitude greater when there is competition than when enzyme concentrations vary independently. If there is positive (C) or negative (E) co-regulation, RNV size is inversely related to |*B*|. When there is competition with positive (D) or negative (F) co-regulation, RNV size is linearly related to |1/*B − e*^0^|. The pseudo-activities **A** were randomly drawn between 0 and 1 000. Initial concentrations **E^0^** were randomly drawn between 0 and 100, **E^0^** was then calibrated to have 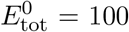. When there is co-regulation, inverses of *B_i_*’s were randomly drawn between 0 and 100 (and randomly multiplied by +1 or − 1 for negative co-regulation), then calibrated to have a sum equal to 1.

##### 3.2.2.2 Case 2: Competition

We did not find any analytical method to study the RNV size near the equilibrium. Graphically, we observed that an enzyme with low pseudo-activity needs larger mutations to get out of the neutral zone (Figure S1B). RNVs were computed numerically from equation 22. We found that RNVs were *inversely* related to enzyme pseudo-activities, unlike in case 1, and hence were positively related to *e*\* (Figure 6B). Another striking observation is that, for a given total enzyme concentration, RNVs are one order of magnitude greater in the case of competition than when there is independence.

##### 3.2.2.3 Case 3: Co-regulation

From equation 28, the relation between *absolute* enzyme concentration and the driving variable *τ* when there is co-regulation can be written as (Supporting Information SI.B.8.1):

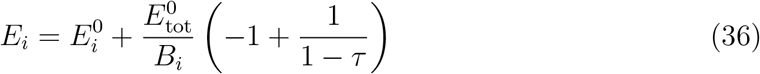

If we consider the superior and inferior limits of the RNV for enzyme *i*, respectively 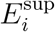 and 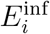 (*i.e.* 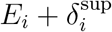 and 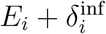), we can write:

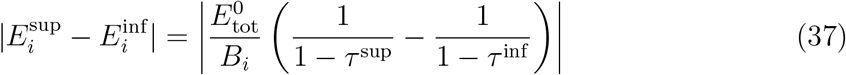

where *τ*^sup^ and *τ*^inf^ are the superior and inferior limits of the driving variable.

We see that RNVs are inversely related to |*B_i_*|’s: the enzymes that have the smallest effect on total enzyme concentration *via* co-regulation have the largest RNV (Figures 6C and 6E).

##### 3.2.2.4 Case 4: Competition plus co-regulation

The same method as previously was applied when we considered jointly competition and co-regulation. From equation 28, we have (Supporting Information SI.B.8.2):

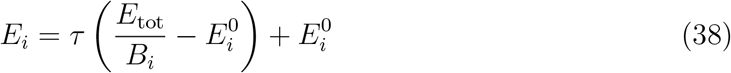

Considering the superior and inferior limits of the RNV for enzyme *i* (see above), we get:

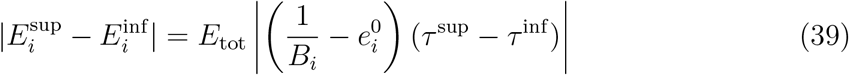

Thus, there is a positive relationship between RNV and 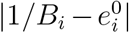, *i.e.* the difference between the theoretical equilibrium (1/*B_i_*) and the initial relative concentration 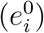 (Figures 6D, 6F), meaning that an enzyme whose concentration is initially far from its theoretical equilibrium will have a large RNV. A geometric representation of this relationship is shown in Figure 7 when there is positive co-regulation.

**Figure 7.**
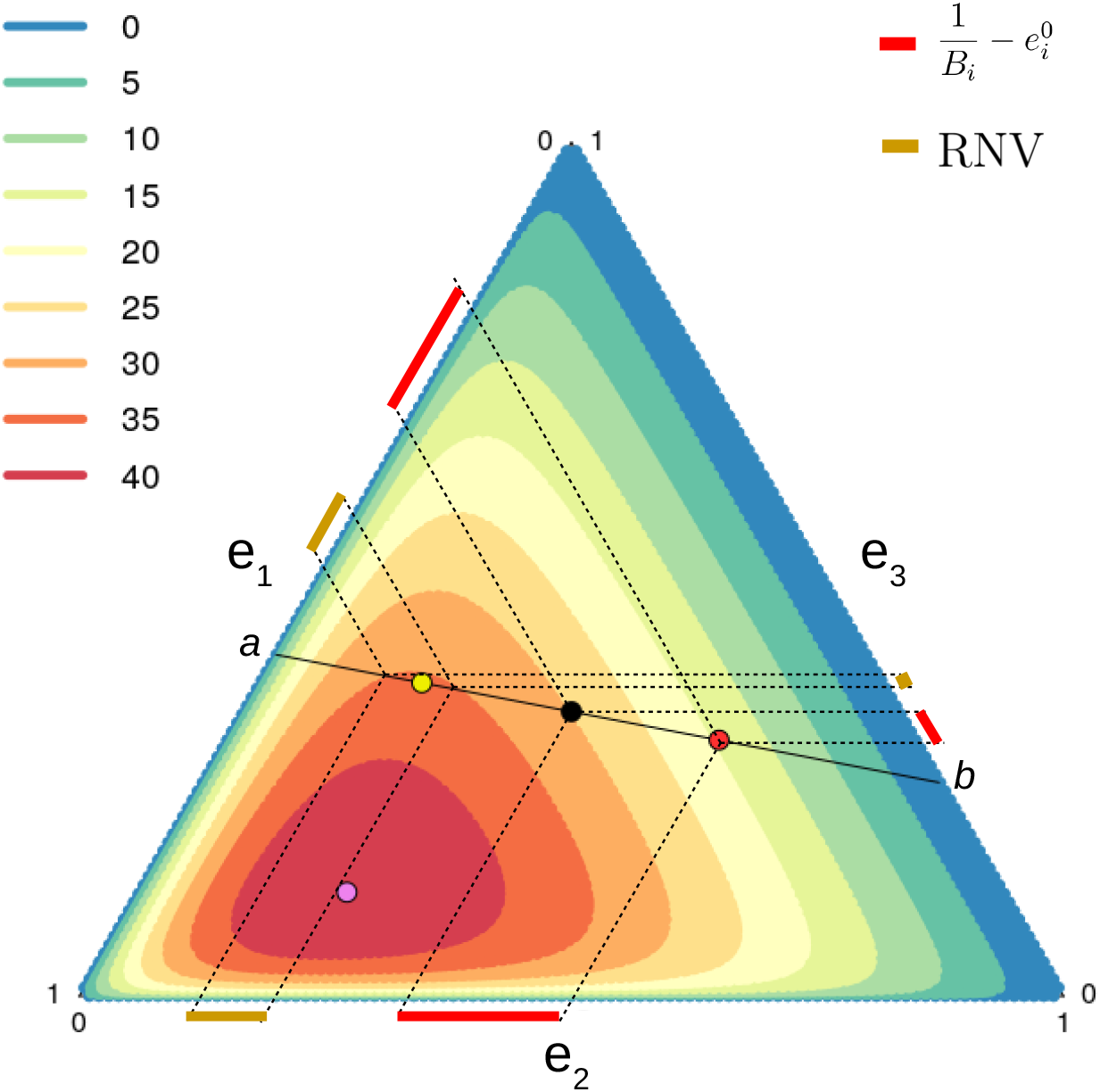
Geometric representation of the RNV when there is both competition and positive co-regulation. The triangle is a top view of the dome of Figure 4A. The color scale represents the flux levels. *a* and *b* are the ends of the line *ε*. Purple point: top of the dome. Black point: initial enzyme concentrations **e^0^**. Yellow point: effective equilibrium 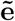. Red point: theoretical equilibrium **e**\*. The red bars outside the triangle represent the difference 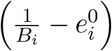 for each of the three enzymes, and the gold bars represent the range of neutral variation (RNV). Note the positive relationship between 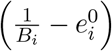 and RNV. The parameters values are as in Figure 4.

Table 3 and Figure 6 summarize the factors determining the ranking order of RNVs depending on the type of constraint.

**Table 3.**
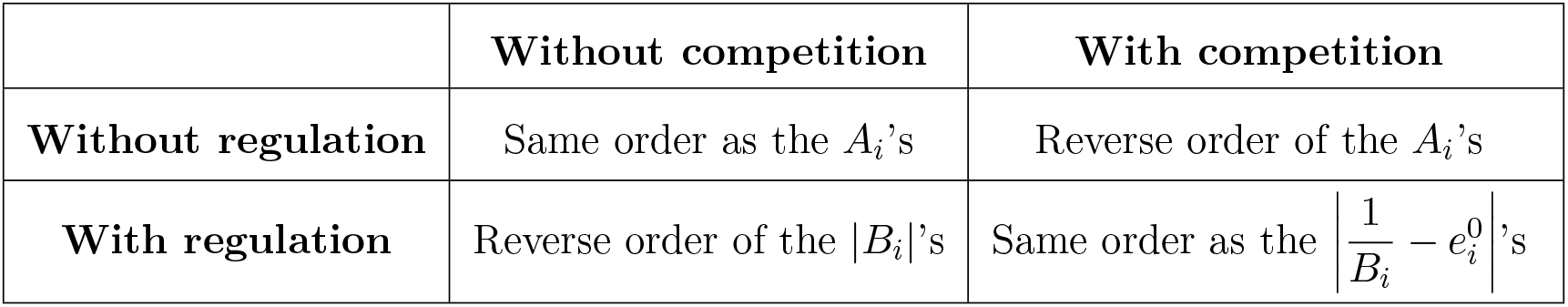
Factors determining the ranking order of RNVs.

#### 3.2.3 Measuring RNVs from simulations of long-term evolution

We tested these theoretical predictions by simulating the long-term evolution of a three-enzyme pathway. For each enzyme and each situation, we computed the RNV limits at each time step between 60,000 and 120,000 generations (*i.e.* once evolutionary equilibrium is reached), then we computed the mean size of the RNVs. The relationship between mean RNV value and *A_i_*, *B_i_* or 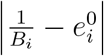 was fully consistent with theoretical predictions (Figure S2).

In addition, the simulations led to two observations.

- When the flux is not limited (independence and positive co-regulation), RNV size increases over the course of generations. For example, if **E^0^ ≈** (21.9, 30.7, 47.3), mean RNV size for enzyme 1 is ≈ 0.13 between 60,000 and 80,000 generations, and ≈ 0.17 between 100,000 and 120,000 generations. The reason is that, over the course of generations, a mutation of a given effect has increasingly less effect on the flux, and therefore on the selection coefficient, because enzyme concentrations increase continuously.
- When there is competition and/or negative co-regulation, the dome-shape of the flux-enzyme relationship can lead to a split of the RNVs. Near the maximum flux, an upper limit of the RNV cannot be defined because mutations are either neutral or deleterious. When the flux moves away from the maximum, advantageous mutations may occur, with the RNV having an upper and a lower limit. But interestingly, another RNV appears on the other side of the dome, due to large mutations that result in flux values that are similar to those caused by mutations of small effect. (Figure S3). The farther the flux is from the maximum, the smaller the two moieties of the RNVs and the stronger the selection to bring the concentrations towards effective equilibrium.

## 4 Discussion

Metabolic fluxes are commonly used as model traits in quantitative genetics and evolutionary studies, benefitting from the development of elaborate theoretical tools that enable to relate “genotype” (the genetically controlled enzyme properties) to phenotype (the flux). The multidimensional flux-enzyme relationship can be viewed as an adaptive landscape and used to analyze the dynamics of enzyme evolution. In this study, we developed a system of differential equations that describes the evolution of enzyme concentrations in a pathway under directional selection for increased metabolic flux. We considered two realistic constraints, separately or jointly, that prevent enzyme concentrations from varying independently from each other, namely competition for cellular resources and co-regulation between enzymes. We defined two types of evolutionary equilibria of relative enzyme concentrations: a theoretical equilibrium and an effective equilibrium, the latter being reached when the flux is limited by certain constraint(s) (tables 1 and 2). In both constrained and unconstrained conditions, mutations become near neutral near the evolutionary equilibrium, however the range of neutral variation (RNVs) of enzyme concentrations differs to a large extent depending on the enzyme properties and the constraints applied (table 3). Simulation results of long-term evolution were fully consistent with the analytical predictions.

### 4.1 Modeling rationale

The effect of an enzyme *j* on the flux of a linear pathway depends on three factors: the ratio of kinetic parameters 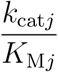, the abundance *E_j_* and the product of the equilibrium constants from the first to the (*j* − 1)^th^ reaction, *K*_1,*j*−1_. Both *E_j_* and 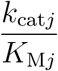 can be genetically variable. Recently, Heckmann et al. (2018) simulated the evolution of turnover numbers *k*_cat*j*_ and showed that their values can hardly be maximized.

In our model, we distinguished the concentration *E_j_* from the “pseudo-activity” 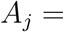 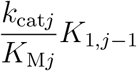 of enzyme *j*, the latter being a systemic property as its value depends on upstream reactions in the pathway. We focused on enzyme concentrations, which display two features that are relevant from an evolutionary point of view:

i. The range of natural variability is expected to be noticeably larger for enzyme concentration than for catalytic efficiency 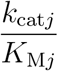. While genetically controlled changes in kinetic parameter values are caused by substitutions at certain sites of the enzyme-coding gene, changes in protein expression can be caused by mutations at multiple *cis* and *trans* elements affecting the gene regulatory network, the translation machinery, mechanisms of protein turnover and stability, etc. (Landry et al. 2007; Zheng et al. 2011; Gruber et al. 2012). This is consistent with the highly polygenic nature of proteomic variations observed in various species (Damerval et al. 1994; Albert and Kruglyak 2015; Chick et al. 2016). Therefore, proteome variability could play a much larger role in the evolution of pathway efficiency that the variability of kinetic parameters. Thus, in our model, we chose to consider mutations that target enzyme concentrations *E_j_*, and let pseudo-activities 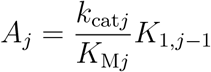 be invariable. Note that kinetic parameters and concentration play an identical role in the expression of the flux, but the constraints applied to *E* do not make sense for *k*_cat_ and *K*_M_.
ii. Enzyme synthesis and degradation are energetically costly for the cell; furthermore, space and ribosome availability are limited (Koehn 1991; de Vienne et al. 2001b; Klumpp et al. 2019) Therefore, protein over-expression can result in protein burden effects (Snoep et al. 1995; Eguchi et al. 2018). Continuous selective pressure for optimizing cellular resource utilization imposes a constraint on total enzyme abundance and a trade-off between enzymes (Albertin et al. 2013).

In addition to the constraint on *E*_tot_, we also considered the possibility that enzyme concentrations were co-regulated. In the cell, multiple mechanisms can result, directly or indirectly, in dependencies between enzyme quantities, *e.g.* operons in prokaryotes, metabolic gene clusters in eukaryotes (Lee and Sonnhammer 2003; Wong and Wolfe 2005) regulation cascades (Albert and Kruglyak 2015), *trans* effects due to transcription factors (Guerin et al. 2016; Lambert et al. 2018), transposable elements (Cowley and Oakey 2013) and microRNAs (Wang et al. 2011), etc. To model the evolutionary implications of these dependencies within a single theoretical framework, we considered that a mutation targeting the concentration of an enzyme modifies the concentrations of the other enzymes. To keep things relatively simple, we chose linear and constant co-regulation coefficients, but we assumed that they could take on any value. When coordinated enzyme expression is due to a common *trans* factor or physically clustered genes, the system of differential equations has to be modified slightly because in this case the mutations target the driving variable *τ* (Supporting Information SI.B.9). This changes the distribution of the mutation effects on the co-regulated enzymes, but does not modify evolutionary equilibria, which do not depend neither on the effects nor on the targets of the mutations. Further developments could rely on more complex models of gene regulatory network functioning and evolution (De Jong 2002; Spirov and Holloway 2013).

The two-step process we used to formalize the mutation effects under competition is not simply theoretically convenient, but may reflect actual events at the molecular level. First, a mutation with a canonical effect *ν* affects the transcription level of an enzyme-coding gene. Second, transcripts are translated proportionally to their abundance up to the maximum total enzyme concentration allowed by the cell. This results in an actual mutation effect on the concentration of the target enzyme and in pseudo-mutation effects on the other enzymes. The process is similar if there is co-regulation in addition to competition.

As in Serohijos et al. (2012) and Heckmann et al. (2018), our evolution model is based on an adaptive dynamics approach that relies on classical population genetics concepts, such as the selection coefficient *s* and the fixation probability *P_f_*, with the assumption of constant population size *N*. Other assumptions were made to render tractable the mathematical developments: small mutation effect relative to enzyme concentration, no fixation of deleterious mutations and large population size. We relaxed these assumptions in the simulations, and this did not affect the predictions of evolutionary equilibria of relative enzyme concentrations and maximum flux values (if applicable).

### 4.2 Constraints shape the adaptive landscape

#### 4.2.1 Optimal enzyme concentrations depend on the type of constraints

The evolutionary equilibria of relative enzyme concentrations depend on the constraints applied. If enzyme concentrations vary independently, there is an inverse relationship between relative concentrations and pseudo-activities **A** at equilibrium, which is intuitively consistent, and both enzyme concentrations and flux can increase indefinitely. This is not biologically realistic because enzyme concentrations are necessarily bounded by limited cellular resources and various regulatory mechanisms. Such constraints shape the adaptive landscape.

When enzymes compete for resources, the adaptive landscape is a dome in the multidimensional space of relative enzyme concentrations; its shape is determined by the enzyme pseudo-activities and its height is determined by the total enzyme concentration. The top of the dome corresponds to the maximum flux reached at equilibrium (Figure 2A). At that point, the relative enzyme concentrations are inversely related to the pseudo-activities **A**, as when enzyme concentrations vary independently, but they are more dispersed. At present, we have no explanation for this result.

Does selection for high flux result in the optimal distribution of enzyme concentrations when there is competition? The theory of optimality of cost minimization has been developed in detail (Klipp and Heinrich 1999; Liebermeister et al. 2004; Noor et al. 2016). Klipp and Heinrich (1999) addressed the question of the optimal distribution of enzyme concentrations in a pathway when there is competition between enzymes. They considered a fixed flux and searched for the distribution that minimizes the total enzyme concentration, which amounts to maximizing the flux with a fixed total concentration, as we did here. Using the Lagrange multiplier method, they obtained the same equilibrium of relative enzyme concentrations as the one we obtained using an adaptive dynamics approach. Thus, the mutation-selection process for increased flux amounts to optimally minimizing the total enzyme concentration allocated to the pathway.

Co-regulation between enzymes prevents reaching this optimum. With co-regulation, the evolutionary trajectories of the relative enzyme concentrations become linear; the orientation of this line is determined by the initial enzyme concentrations and the co-regulation coefficients. Hence the flux follows the curve defined by the intersection of the dome and the upright plane drawn from the line (Figure 4). The maximal flux value on this curve, 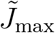, is a local maximum corresponding to the effective equilibrium of relative enzyme concentrations.

If there is co-regulation, with or without competition, the theoretical equilibrium of relative enzyme concentrations depends on the co-regulation coefficients, according to the simple relation 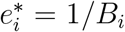, where the global regulation coefficient *B_i_* quantifies the effect of *E_i_* variation on the total enzyme concentration. This evolutionary equilibrium implies that:

i. The higher the effect of enzyme concentration variation on the total concentration, the lower the relative concentration of this enzyme at evolutionary equilibrium;
ii. Pseudo-activities *A_i_* do not appear in this relation. This does not mean that the kinetic parameters and equilibrium constants do not affect the evolutionary trajectory when there is co-regulation. Indeed, *A_i_*’s determine the shape of the adaptive landscape, and hence the flux value at evolutionary equilibrium;
iii. This equilibrium is an evolutionary black hole: as one gets closer, the effect of mutations on *relative* enzyme concentrations decreases, and is cancelled out at this equilibrium irrespective of the effects of these mutations. This property has an important consequence when there is competition, because it can prevent the flux from reaching its local maximum 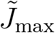. If the initial flux value is lower than the flux value at theoretical equilibrium *and* both values are on the same side of the curve (*i.e.* the derivatives at these points have the same sign), selection will be unable to bring the flux value above the theoretical equilibrium (Figures 4B and 5A);
iv. If there is negative co-regulation, at least one *B_i_* is negative and theoretical equilibrium is not achievable. The flux will then reach a maximum corresponding to the effective equilibrium, whether there is competition or not.

In summary, co-regulation can be evolutionarily costly, since it can prevent the flux from reaching the maximum value given the total enzyme concentration allocated to the pathway. Therefore, increasing fitness by increasing the flux should imply having mutations that modify the co-regulation coefficients, *i.e.* that modify the orientation of line *ε* to bring 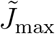 closer to *J*_max_.

#### 4.2.2 The Range of Neutral Variation of enzyme concentrations depends markedly on enzyme properties, constraints and/or initial enzyme concentrations

From visual inspection of the simulation results, we found that relative enzyme concentrations fluctuate close to their evolutionary equilibrium, which is consistent with the mutation-selection-drift balance: selection drives the system towards equilibrium, but near the equilibrium, mutations become nearly neutral and genetic drift dominates (Figures 2B, 3C, 5A, 5C). Near the point of evolutionary equilibrium, mutations have a negligible effect on the flux, and therefore are in the neutral zone. We compared analytically the range of variation of enzyme concentrations in the neutral zone (RNVs) at evolutionary equilibrium under different conditions.

When enzyme concentrations vary independently, the RNVs are positively related to the pseudo-activities; the opposite is found when there is competition (Figures 6A and 6B). Thus, competition results in a positive relationship between RNVs and enzyme concentrations. In addition, competition strongly increases all RNVs for a given *E*_tot_.

When there is only co-regulation, enzymes whose variation has the largest effect on total concentration – hence the smallest effective equilibrium of relative enzyme concentrations – have the smallest RNVs. With both competition and co-regulation, the more an enzyme is initially far from the effective equilibrium, the larger its RNV (Figure 7). Therefore, co-regulation abolishes the effect of *A* on RNVs, whether there is competition or not, as they did for the theoretical equilibrium values.

Interestingly, the existence of a flux maximum can lead to a discontinuous RNV. One part of the RNV corresponds to mutations of small effect, another part to mutations of large effect; the latter push the enzyme concentrations beyond the top of the dome, where flux values are similar to the resident flux. This optimum overshoot is inherent to Fisher’s geometric model of adaptive landscapes (Orr 1998), and is more common when phenotypes are close to the optimum (Scott and Queller 2019).

The magnitude of the RNV should be reflected in the level of polymorphism of the genes affecting enzyme abundance, because high-RNV enzymes can accumulate more neutral mutations in these genes than low-RNV enzymes. It would be informative to compare the levels of variability of enzyme abundances within a pathway and investigate how these are related to kinetic parameters and/or known co-regulatory interactions. Quantitative proteomics databases could be exploited for this purpose.

### 4.3 Conclusion and future prospects

We combined MCT principles, an adaptive dynamics approach and simulations to analyze the evolution of enzyme concentrations in metabolic pathways under selection for increased flux. We showed that the adaptive landscape is shaped by enzyme properties and by constraints applied to the system. When there is both competition for cellular resources and co-regulation that limits the flux, which is likely to be the most common situation in a cell, selective pressures lead to equilibria of enzyme concentrations and to ranges of neutral variation that not only depend on enzyme properties and the type of constraints but also on the enzyme concentrations at the start of the selective process. This evolutionary contingency comes from the effect of co-regulation between enzymes and could be mitigated by mutations modifying certain regulatory parameters. In addition, the assumption that all enzymes of the pathway are co-regulated can be relaxed by considering groups of differently co-regulated enzymes (manuscript in preparation).

## Acknowledgements

We warmly thank Dr. Wolfram Liebermeister and Dr. Arnaud Le Rouzic for their careful reading of the manuscript and their very helpful suggestions. We also thank Pr. Amandine Veber for her useful suggestions regarding the differential equation system of enzyme evolution and Hélène Citerne for language corrections of the manuscript.

CC was supported by a PhD thesis grant from the French Ministère de l’Enseignement Supérieur, de la Recherche et de l’Innovation.

The authors declare no conflict of interest.

## Author contributions

CD and DdV initiated the project. CD, MLL, GT, CC and DdV developed the model. GT, MLL and CC performed the analysis. CD and MLL wrote the simulation code. CC developed the dedicated R package. CC and DdV wrote the manuscript, with the contribution of CD.

## Appendix A Glossary of mathematical symbols

### Latin symbols

*A_i_*: Pseudo-activity of enzyme *i* (the parameter includes the kinetic properties of the enzyme and thermodynamic equilibrium of upstream reactions)
A: Vector of pseudo-activities
*B_i_*: Global co-regulation coefficient, representing effect of a mutation of *E_i_* on *E*_tot_ due to co-regulation
*E_i_*: Concentration of enzyme *i*
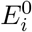: Initial concentration of enzyme *i*
E: Vector of enzyme concentrations, considered as the genotype
E^0^: Vector of initial enzyme concentrations
**E**^(**1**)^: Vector of enzyme concentrations after a mutation
**E**^(**2**)^: Vector of enzyme concentrations after a mutation and the application of a constraint on *E*_tot_
E^m^: Vector of enzyme concentrations of the mutant
E^r^: Vector of enzyme concentrations of the resident
**E**^(*t*)^: Vector of concentrations at time *t*
*E*_tot_: Total enzyme concentration in the metabolic pathway
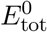: Initial total enzyme concentration
*e_i_*: Relative concentration of enzyme *i*
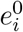: Initial relative concentration of enzyme *i*
e: Vector of relative enzyme concentrations
e^0^: Vector of initial relative enzyme concentrations
**e**\*: Theoretical equilibrium of relative enzyme concentrations
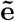: Effective equilibrium of relative enzyme concentrations, corresponding to a maximum of the flux
*ε*: Line in the space of relative enzyme concentrations along which the relative concentrations vary in the case of co-regulation
*J*: Metabolic flux, considered as a phenotype that is proportional to fitness
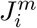: Flux resulting from a mutation targeting enzyme *i*
*J^r^*: Flux of the resident
*K_i_*: Equilibrium constant of reaction *i*, catalyzed by enzyme *i*
*K*_1,*i*_: Product of the equilibrium constants from the first to the *i*^th^ reaction: 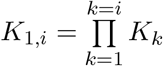
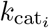: Catalytic constant of enzyme *i*
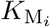: Michaelis-Menten constant of enzyme *i* for its substrate
**M**_*β*_: Matrix of co-regulation coefficients
*N*: Effective population size
*n*: Number of enzymes in the metabolic pathway
*P_f_*: Fixation probability of a mutation
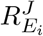: Flux response coefficient of enzyme *i*
S_*i*_: Metabolite *i*, product of reaction *i* and substrate of reaction *i* + 1
*s_i_*: Selection coefficient of a mutation targeting enzyme *i*
*t*: Evolutionary time
*w*: Fitness
*w^m^*: Mutant fitness
*w^r^*: Resident fitness
*X*: Constant representing the environment: *X* = *X*_0_ − *X_n_*/ *K*_1,*n*_
*X*_0_: Concentration of the input substrate of the pathway
*X_n_*: Concentration of the output product of the pathway

### Greek symbols

*α_ij_*: Redistribution coefficient, expressing the variation of *E_j_* due to mutation affecting enzyme *i*
*β_ij_*: Co-regulation coefficient of enzyme *i* on *j*
*γ_ij_*: Competition coefficient, expression of *α_ij_* in the case of competition alone
*δ_i_*: Actual effect of a mutation affecting enzyme *i*
*μ*: Mutation rate at each time unit
*ν*: Canonical effect of a mutation
*τ*: Driving variable of the line *ε* in the space of relative enzyme concentrations in the case of co-regulation.

## Notes

### Competing Interest Statement

The authors have declared no competing interest.

