## Supporting Information for "Evolution of enzyme levels in metabolic pathways: A theoretical approach"

Charlotte Coton, Grégoire Talbot, Maud Le Louarn, Christine Dillmann, Dominique de Vienne

May 4, 2021

### SI Supporting Information

#### SI.A Supplementary Figures

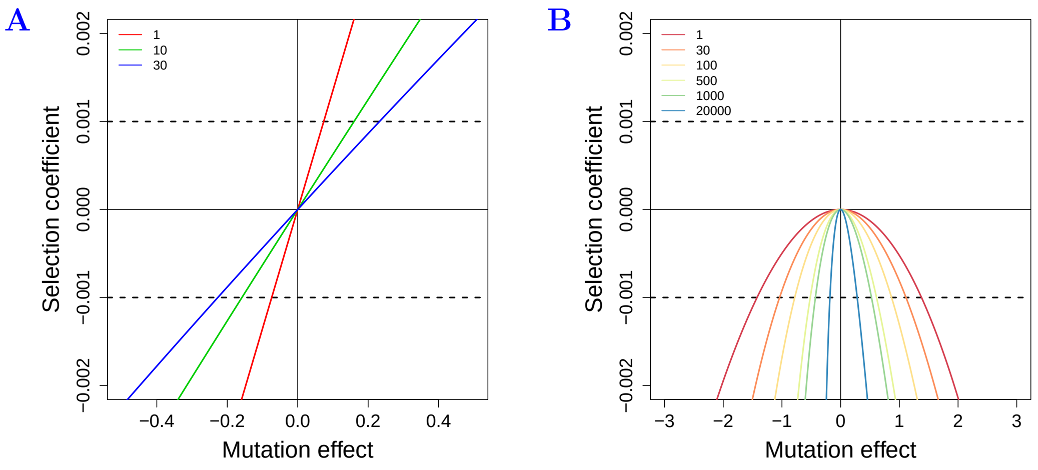

**Figure S1. Range of neutral variation in the case of independence and competition.** (A) Relationship between the selection coefficient $s_i = \Delta J/J$ and the actual effect of a mutation $\delta_i$ of enzyme $i$ at evolutionary equilibrium of relative enzyme concentrations in a pathway of three enzymes that vary independently. The pseudo-activities are $A_1 = 1$ (red), $A_2 = 10$ (green) and $A_3 = 30$ (blue). $E_i^0$'s are respectively 55.99, 25.99 and 18.02, corresponding to the equilibrium values when $E_{\text{tot}}^0 = 100$. The neutral zone is located between the two horizontal dotted lines at $s_i = \pm 0.001$. For the most active enzyme (blue curve), larger mutation effects are required to get out the neutral zone than for the least active enzyme (red curve). The curve of enzyme with intermediate pseudo-activity (green curve) is between that of the most and least active enzyme. Therefore there is a positive relationship between pseudo-activity and RNV size. (B) Representation as in (A) of a six-enzyme pathway with competition ($E_{\text{tot}} = 100$). The pseudo-activities are: $A_1 = 10$, $A_2 = 30$, $A_3 = 100$, $A_4 = 500$, $A_5 = 1\ 000$ and $A_6 = 20\ 000$ (color palette from red to blue). The selection coefficients are all negative because any deviation from the equilibrium value results in negative $s_i$. Unlike in (A) the most active enzymes have the narrowest range of neutral variation.

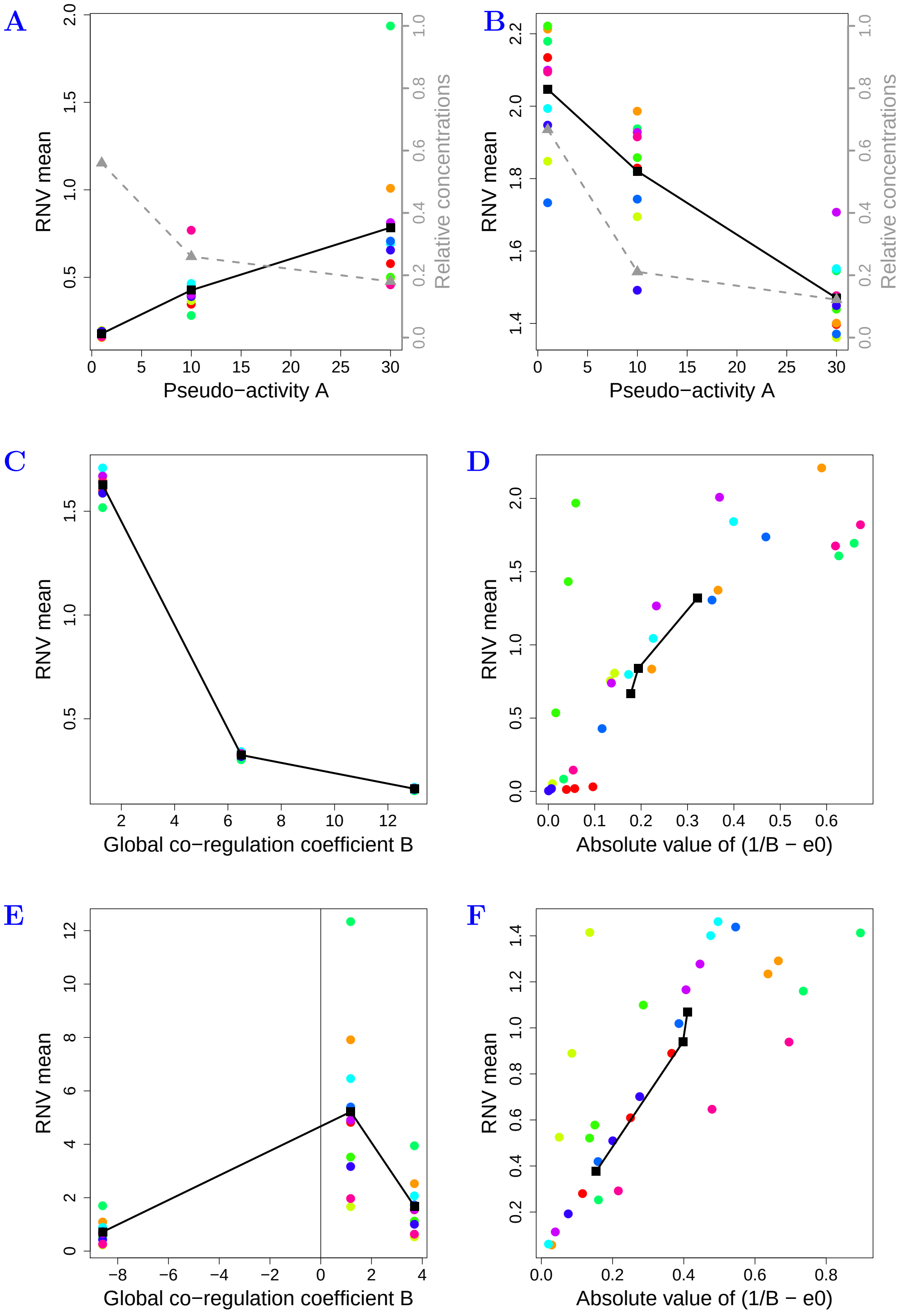

**Figure S2. Relationship between constraint-dependent factors and RNV from simulations of long-term evolution of a three-enzyme pathway.** Representation is as in Figure 6. For each of the ten simulations (one color *per* simulation), the RNV of each enzyme was computed as the mean of the RNV sizes between 60,000 and 120,000 generations, *i.e.* once equilibrium is reached. Black points are the mean values of all simulations. Gray points and dashed line in (A) and (B) indicate the relative enzyme concentrations at evolutionary equilibrium. The colored points in (D) and (F) are not vertically aligned because the different simulations do not have the same $e^0$ values. We see that the results are qualitatively the same as in Figure 6. Note the positive relationship between RNV size and RNV variability. Parameter values are as in Figures 1 to 5.

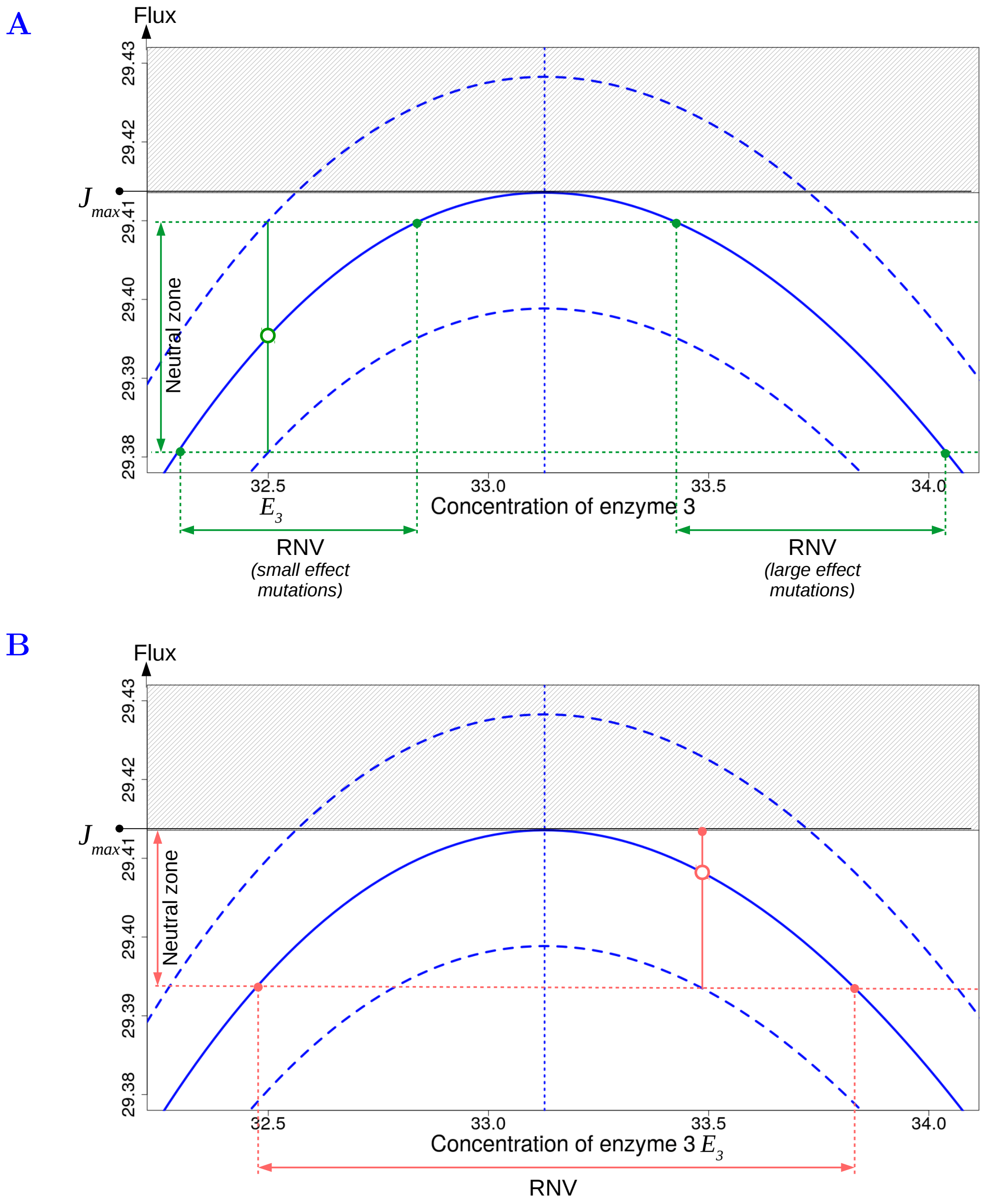

**Figure S3. Range of neutral variation (RNV) in the case of negative co-regulation.** The flux (blue curve) is function of the concentration of one enzyme (enzyme 3) in a three-enzyme pathway. $\tilde{J}_{\max}$ is reached at effective equilibrium $\tilde{E}_3 = 33.13$ (vertical blue dotted line). The upper and lower dashed blue curves delimit the neutral zone of flux variation. Due to the existence of a maximum flux, the gray area is unattainable. (A) The concentration of enzyme 3 is far from the equilibrium, at $E_3 \approx 32.5$. The vertical green line shows the amplitude of the neutral zone (between the horizontal green dotted lines). The corresponding RNV splits into two parts (horizontal green left and right arrows), because mutations of large effect can lead to flux values that are similar to those caused by mutations of small effect. (B) The concentration of enzyme 3 is closer to the effective equilibrium, at $E_3 \approx 33.5$. Here the upper limit of the neutral zone cannot be achieved because it exceeds the maximum flux. The continuous RNV (horizontal red left and right arrow) is bounded by the enzyme concentrations 32.45 and 33.85 corresponding to the inferior limit of neutral zone on both sides of the curve. Parameter values are: $X = 1$, $A_1 = 1$, $A_2 = 10$, $A_3 = 30$, $E_{\text{tot}} = 100$, $\beta_{12} = -8.75$, $\beta_{23} = -2.25$.

### SI.B Mathematical proofs

#### SI.B.1 Properties of the redistribution coefficient and mutation effects

**SI.B.1.1 The matrix of pairwise co-regulation coefficients** From any vector $\boldsymbol{\beta}$ of $\beta_{i-}$'s, all elements of the matrix $\mathbf{M}_\beta$ of pairwise co-regulation coefficients can be deduced from the relation $\mathbf{M}_\beta = \boldsymbol{\beta}^{\text{inv}}\boldsymbol{\beta}^\mathsf{T}$, where $\boldsymbol{\beta}^{\text{inv}}$ is the vector of inverse elements of $\boldsymbol{\beta}$ (Table S1). Note that $\mathbf{M}_\beta$ is symmetric.

**Table S1. Matrix $\mathbf{M}_\beta$ of the co-regulation coefficients**.

| $i \backslash j$ | $\beta_{-1}$ | $\beta_{-2}$ | $\cdots$ | $\beta_{-i}$ | $\beta_{-j}$ | $\cdots$ | $\beta_{-n}$ | Sum |
|---|---|---|---|---|---|---|---|---|
| $\beta_{1-}$ | 1 | $\beta_{i2}/\beta_{i1}$ | $\cdots$ | $\beta_{ii}/\beta_{i1}$ | $\beta_{ij}/\beta_{i1}$ | $\cdots$ | $\beta_{in}/\beta_{i1}$ | $B_1$ |
| $\beta_{2-}$ | $\beta_{i1}/\beta_{i2}$ | 1 | $\cdots$ | $\beta_{ii}/\beta_{i2}$ | $\beta_{ij}/\beta_{i2}$ | $\cdots$ | $\beta_{in}/\beta_{i2}$ | $B_2$ |
| $\vdots$ | $\vdots$ | $\vdots$ | $\ddots$ | $\cdots$ | $\cdots$ | $\cdots$ | $\cdots$ | $\vdots$ |
| $\beta_{i-}$ | $\beta_{i1}/\beta_{ii}$ | $\beta_{i2}/\beta_{ii}$ | $\vdots$ | 1 | $\beta_{ij}/\beta_{ii}$ | $\cdots$ | $\beta_{in}/\beta_{ii}$ | $B_i$ |
| $\beta_{j-}$ | $\beta_{i1}/\beta_{ij}$ | $\beta_{i2}/\beta_{ij}$ | $\vdots$ | $\beta_{ii}/\beta_{ij}$ | 1 | $\cdots$ | $\beta_{in}/\beta_{ij}$ | $B_j$ |
| $\vdots$ | $\vdots$ | $\vdots$ | $\vdots$ | $\vdots$ | $\vdots$ | $\ddots$ | $\cdots$ | $\vdots$ |
| $\beta_{n-}$ | $\beta_{i1}/\beta_{in}$ | $\beta_{i2}/\beta_{in}$ | $\vdots$ | $\beta_{ii}/\beta_{in}$ | $\beta_{ij}/\beta_{in}$ | $\vdots$ | 1 | $B_n$ |

**SI.B.1.2 Properties of the global co-regulation coefficients** Let a group of co-regulated enzymes, *i.e.* $\beta_{ij} \neq 0$ for all $(i, j)$. Calculating global co-regulation coefficient leads to:

$$\forall i \quad B_i = \sum_{k=1}^{n} \beta_{ik} = \sum_{k=1}^{n} \beta_{ij}\beta_{jk} = \beta_{ij} \sum_{k=1}^{n} \beta_{jk} = \beta_{ij} B_j \tag{S1}$$

$$\boxed{B_i = \beta_{ij} B_j} \tag{S2}$$

Therefore:

1. If $\forall j \neq i$, $\beta_{ij} > 0$, then $B_i = \sum_{k=1}^{n} \beta_{ik} > 1$ and $B_j > 1$.
2. If it exists at least $(i, j)$ such as $\beta_{ij} < 0$ and $B_i > 0$, then $B_j < 0$.
3. If it exists at least $(i, j)$ such as $\beta_{ij} < 0$ and $B_i < 0$, then $B_j > 0$.

Thus, if there is at least one negative co-regulation coefficient, there is at least one negative global co-regulation coefficient.

**SI.B.1.3 Mutation effect and redistribution coefficient in the case of competition** Let us consider a resident population with genotype $\mathbf{E}$ and a mutation of canonical effect $\nu$ targeting enzyme $i$. To formalize the consequences of the mutation on genotype $\mathbf{E}$ when there is competition for cellular resources, *i.e.* constant total enzyme concentration, we considered a two-step process: first, the mutation modifies the concentration of enzyme $i$, second, there is reduction from $E_{\text{tot}} + \nu$ to $E_{\text{tot}}$, with proportional redistribution of enzyme concentrations:

| Concentration | Enzyme $i$ | Enzyme $j \neq i$ | Total concentration |
|---|---|---|---|
| Resident | $E_i$ | $E_j$ | $E_{\text{tot}}$ |
| After mutation targeting $i$ | $E_i^{(1)} = E_i + \nu$ | $E_j^{(1)} = E_j$ | $E_{\text{tot}}^{(1)} = E_{\text{tot}} + \nu$ |
| After constraint on $E_{\text{tot}}$ | $E_i^{(2)} = E_i + \delta_i$ | $E_j^{(2)}$ | $E_{\text{tot}}$ |

Assuming that the proportions of enzyme concentrations are conserved through the redistribution, we have for the targeted enzyme $i$:

$$\frac{E_i + \nu}{E_{\text{tot}} + \nu} = \frac{E_i^{(2)}}{E_{\text{tot}}} \tag{S3}$$

The actual effect of the mutation is:

$$\delta_i = E_i^{(2)} - E_i = E_{\text{tot}} \frac{E_i + \nu}{E_{\text{tot}} + \nu} - E_i$$

As $E_i = e_i E_{\text{tot}}$, we get after rearrangements:

$$\boxed{\delta_i = \frac{1 - e_i}{\frac{1}{\nu} + \frac{1}{E_{\text{tot}}}}} \tag{S4}$$

For enzymes $j \neq i$, we have:

$$\frac{E_j}{E_{\text{tot}} + \nu} = \frac{E_j^{(2)}}{E_{\text{tot}}} \tag{S5}$$

So

$$E_j^{(2)} = \frac{E_j}{E_{\text{tot}} + \nu} E_{\text{tot}}$$

In order to express $E_j^{(2)}$ in terms of $E_j$ and $\delta_i$, we write:

$$E_j^{(2)} - E_j = E_{\text{tot}} \frac{E_j}{E_{\text{tot}} + \nu} - E_j = -\frac{E_j \nu}{E_{\text{tot}} + \nu} = -\frac{e_j}{\frac{1}{\nu} + \frac{1}{E_{\text{tot}}}}$$

and using equation S4, we get:

$$\boxed{E_j^{(2)} = E_j - \frac{e_j}{1 - e_i} \delta_i} \tag{S6}$$

Thus, the term $\dfrac{-e_j}{1 - e_i}$ is the redistribution coefficient $\alpha_{ij}$ when there is competition.

**SI.B.1.4 Mutation effect and redistribution coefficient in the case of competition plus co-regulation** If there is co-regulation in addition to competition, the two-step process described above is slightly modified, because the mutation targeting enzyme $i$ also affects indirectly the co-regulated enzymes. So we introduce the co-regulation coefficients $\beta_{ij}$ in the first step:

| Concentration | Enzyme $i$ | Enzyme $j \neq i$ | Total concentration |
|---|---|---|---|
| Resident | $E_i$ | $E_j$ | $E_{\text{tot}}$ |
| Mutation + co-regulation | $E_i^{(1)} = E_i + \nu$ | $E_j^{(1)} = E_j + \beta_{ij}\nu$ | $E_{\text{tot}}^{(1)} = E_{\text{tot}} + B_i \nu$ |
| Constraint on $E_{\text{tot}}$ | $E_i^{(2)} = E_i + \delta_i$ | $E_j^{(2)}$ | $E_{\text{tot}}$ |

Assuming that the proportions of the enzyme concentrations are conserved through the redistribution, we have for the targeted enzyme $i$:

$$\frac{E_i + \nu}{E_{\text{tot}} + B_i \nu} = \frac{E_i^{(2)}}{E_{\text{tot}}} \tag{S7}$$

The actual effect of the mutation is:

$$\delta_i = E_i^{(2)} - E_i$$

So, from equation S7, we have:

$$\delta_i = E_{\text{tot}} \frac{E_i + \nu}{E_{\text{tot}} + B_i \nu} - E_i$$

As $E_i = e_i E_{\text{tot}}$, we get after rearrangements:

$$\boxed{\delta_i = \frac{1 - B_i e_i}{\frac{1}{\nu} + \frac{B_i}{E_{\text{tot}}}}} \tag{S8}$$

For enzymes $j \neq i$, we have:

$$\frac{E_j + \beta_{ij} \nu}{E_{\text{tot}} + B_i \nu} = \frac{E_j^{(2)}}{E_{\text{tot}}} \tag{S9}$$

So:

$$E_j^{(2)} = \frac{E_j + \beta_{ij} \nu}{E_{\text{tot}} + B_i \nu} E_{\text{tot}}$$

In order to express $E_j^{(2)}$ in terms of $E_j$ and parameters depending on $E_i$, we write:

$$E_j^{(2)} - E_j = E_{\text{tot}} \frac{E_j + \beta_{ij} \nu}{E_{\text{tot}} + B_i \nu} - E_j = \frac{E_{\text{tot}} \beta_{ij} \nu - E_j B_i \nu}{E_{\text{tot}} + B_i \nu} = \frac{\beta_{ij} - B_i e_j}{\frac{1}{\nu} + \frac{B_i}{E_{\text{tot}}}}$$

and using equation S8, we get:

$$\boxed{E_j^{(2)} = E_j + \frac{\beta_{ij} - B_i e_j}{1 - B_i e_i} \delta_i} \tag{S10}$$

Thus, when there is competition plus co-regulation, the redistribution coefficient can be written:

$$\boxed{\alpha_{ij} = \frac{\beta_{ij} - B_i e_j}{1 - B_i e_i}} \tag{S11}$$

From this expression, we can distinguish two particular cases:

1. If there is only competition, *i.e.* $\forall i,\ j \neq i,\ \beta_{ij} = 0$ and $B_i = 1$, we have:

$$\alpha_{ij} = \frac{-e_j}{1 - e_i}$$

2. If all enzymes are co-regulated, we have $\beta_{ij} = B_i / B_j$ (equation S2), and therefore:

$$\alpha_{ij} = \frac{1/B_j - e_j}{1/B_i - e_i}$$

   Note that this relation is also valid for $\alpha_{ii} = 1$ and has the same properties as the $\beta$'s, *i.e.* $\alpha_{ij} = \alpha_{ik} \alpha_{kj}$ and $\alpha_{ij} = 1/\alpha_{ji}$.

In the case of co-regulation without competition, $\mathbf{E^m}$ is equal to $\mathbf{E^{(1)}}$, and we have $\alpha_{ij} = \beta_{ij}$ for all $(i, j)$ and $\delta_i = \nu$.

##### SI.B.2 Explicit expression of the response coefficient

The response coefficient is defined by:

$$\forall i \quad R^J_{E_i} = \frac{\partial J/J}{\partial E_i/E_i}$$

Introducing the expression of $J$ (equation 1), we get:

$$\forall i \quad R^J_{E_i} = -X\frac{E_i}{J}\frac{\frac{\partial \sum\limits_{j=1}^{n}\frac{1}{A_jE_j}}{\partial E_i}}{\left(\sum\limits_{j=1}^{n}\frac{1}{A_jE_j}\right)^2}$$

$$= X\frac{E_i}{\frac{X}{\sum\limits_{j=1}^{n}\frac{1}{A_jE_j}}}\frac{\sum\limits_{j=1}^{n}\frac{\partial E_j/\partial E_i}{A_jE_j^2}}{\left(\sum\limits_{j=1}^{n}\frac{1}{A_jE_j}\right)^2}$$

$$= E_i\frac{\sum\limits_{j=1}^{n}\frac{\partial E_j/\partial E_i}{A_jE_j^2}}{\sum\limits_{j=1}^{n}\frac{1}{A_jE_j}}$$

As $\alpha_{ij}$ is the derivative of $E_j$ with respect to $E_i$ *(Material & Methods)*, we obtain:

$$\boxed{\forall i, \quad R^J_{E_i} = E_i\frac{\sum\limits_{j=1}^{n}\frac{\alpha_{ij}}{A_jE_j^2}}{\sum\limits_{j=1}^{n}\frac{1}{A_jE_j}} = e_i\frac{\sum\limits_{j=1}^{n}\frac{\alpha_{ij}}{A_je_j^2}}{\sum\limits_{j=1}^{n}\frac{1}{A_je_j}}} \tag{S12}$$

##### SI.B.3 Selective neutrality and range of neutral variation (RNV)

**SI.B.3.1 Expressing the selection coefficient in term of the response coefficient** Let us consider a mutation of actual effect $\delta_i$ that changes the concentration of enzyme $i$ of the resident from $E_i^r$ to $E_i^m$:

$$\delta_i = E_i^m - E_i^r = \Delta E_i$$

and the flux from $J^r$ to $J^m$:

$$J^m - J^r = \Delta J$$

Because the flux is assumed to be proportional to fitness, the coefficient of selection of the mutant is:

$$s_i = \left(\frac{\Delta J}{J}\right)_i$$

If the mutation effect $\delta_i$ is small compared to concentration $E_i$, we have:

$$\frac{s_i}{\delta_i/E_i} = \frac{\Delta J/J}{\Delta E_i/E_i} \approx \frac{\partial J/J}{\partial E_i/E_i} \tag{S13}$$

where we recognize the response coefficient of enzyme $i$. Therefore, the coefficient of selection is:

$$\boxed{s_i \approx R^J_{E_i}\frac{\delta_i}{E_i}} \tag{S14}$$

**SI.B.3.2 Computing the limits of the RNV** To determine the values $\delta_i^{\mathrm{sup}}$ and $\delta_i^{\mathrm{inf}}$ that respectively define the superior and inferior limits of the RNV, we used the equation of the selection coefficient that depends on $J^r$ and $J_i^m$, the flux value before and after a mutation targeting the enzyme $i$ (equation 15).

Writing

$$J_i^m = J^r(s_i + 1)$$

and replacing the $J$'s by their respective expressions, we get:

$$\frac{X}{\sum_{j=1}^{n} \frac{1}{A_j(E_j + \alpha_{ij}\delta_i)}} = (s_i + 1)\frac{X}{\sum_{j=1}^{n} \frac{1}{A_j E_j}}$$

Taking the inverse, we obtain:

$$\boxed{\sum_{j=1}^{n} \frac{1}{A_j(E_j + \alpha_{ij}\delta_i)} = \frac{1}{s_i + 1}\sum_{j=1}^{n} \frac{1}{A_j E_j}} \qquad \text{(S15)}$$

Resolving this equation for $s_i = 1/N$ (resp. $s_i = -1/N$) gives $\delta_i^{\mathrm{sup}}$ (resp. $\delta_i^{\mathrm{inf}}$).

##### SI.B.4 Evolutionary equilibrium in the case of independence (case 1)

When there is independence between enzyme concentrations, we have $\alpha_{ii} = 1$ for all $i$ and $\alpha_{ij} = 0$ for all $j \neq i$ and the differential equation system describing the evolution of the absolute concentrations becomes:

$$\forall j \quad \frac{\partial E_j}{\partial t} = 2\mu s_j \delta_j$$

and the differential equation for total concentration writes:

$$\begin{aligned}\frac{\partial E_{\mathrm{tot}}}{\partial t} &= \sum_{k=1}^{n} \frac{\partial E_k}{\partial t} \\ &= \sum_{k=1}^{n} 2\mu s_k \delta_k \\ &= 2\mu \sum_{k=1}^{n} s_k \delta_k\end{aligned}$$

These expressions allow us to write the differential equation system describing the evolution of the *relative* concentrations:

$$\forall j \quad \frac{\partial e_j}{\partial t} = \frac{\partial (E_j/E_{\mathrm{tot}})}{\partial t} = \frac{\frac{\partial E_j}{\partial t}E_{\mathrm{tot}} - E_j\frac{\partial E_{\mathrm{tot}}}{\partial t}}{(E_{\mathrm{tot}})^2}$$

By replacing the derivatives by their expression, we get:

$$\begin{aligned}\forall j \quad \frac{\partial e_j}{\partial t} &= \frac{E_{\mathrm{tot}} 2\mu s_j \delta_j - e_j E_{\mathrm{tot}} 2\mu \sum_{k=1}^{n} s_k \delta_k}{(E_{\mathrm{tot}})^2} \\ &= \frac{2\mu}{E_{\mathrm{tot}}}(s_j \delta_j - e_j \sum_{k=1}^{n} s_k \delta_k)\end{aligned}$$

Due to the relationship between selection and response coefficients (equation S14), we can write:

$$\forall j \quad \frac{\partial e_j}{\partial t} = \frac{2\mu}{E_{\text{tot}}}(R^J_{E_j}\frac{\delta_j^2}{E_j} - e_j\sum_{k=1}^{n} R^J_{E_k}\frac{\delta_k^2}{E_k}) = \frac{2\mu}{E_{\text{tot}}^2}(R^J_{E_j}\frac{\delta_j^2}{e_j} - e_j\sum_{k=1}^{n} R^J_{E_k}\frac{\delta_k^2}{e_k})$$

As the steady state of relative enzyme concentrations is defined by

$$\forall j, \quad \frac{\partial e_j}{\partial t} = 0$$

the relative concentrations $e_j^*$ at evolutionary equilibrium can be found by solving:

$$\forall j, \quad R^{J*}_{E_j}\frac{\delta_j^2}{e_j^*} = e_j^*\sum_{k=1}^{n} R^{J*}_{E_k}\frac{\delta_k^2}{e_k^*}$$

When there is independence, the actual and canonical effects of the mutation are equal, *i.e.* $\delta_i = \nu$ for all $i$. So we have:

$$\forall j, \quad R^{J*}_{E_j}\frac{\nu^2}{e_j^*} = \nu^2 e_j^*\sum_{k=1}^{n} \frac{R^{J*}_{E_k}}{e_k^*},$$

Leading to:

$$\forall j \quad e_j^* = \frac{\frac{R^{J*}_{E_j}}{e_j^*}}{\sum_{k=1}^{n}\frac{R^{J*}_{E_k}}{e_k^*}}$$

Replacing the response coefficient by its expression in the case of independence (equation S12), *i.e.* with $\alpha_{ii} = 1$ and $\alpha_{ij} = 0, \forall j \neq i$, we obtain:

$$\forall j, \quad e_j^* = \frac{\frac{\frac{1}{A_j e_j^* E_{\text{tot}}}}{\sum_{l=1}^{n}\frac{1}{A_l E_l}} \Big/ e_j^*}{\sum_{k=1}^{n}\left(\frac{\frac{1}{A_k e_k^* E_{\text{tot}}}}{\sum_{l=1}^{n}\frac{1}{A_l E_l}} \Big/ e_k^*\right)} = \frac{\frac{1}{A_j (e_j^*)^2}}{\sum_{k=1}^{n}\frac{1}{A_k (e_k^*)^2}}$$

Then:

$$\sum_{k=1}^{n}\frac{1}{A_k (e_k^*)^2} = \frac{1}{A_j (e_j^*)^3}$$

As $\sum_{k=1}^{n}\frac{1}{A_k (e_k^*)^2}$ is constant, this equality is valid for any enzyme:

$$\forall i, \ \forall j \quad \sum_{k=1}^{n}\frac{1}{A_k (e_k^*)^2} = \frac{1}{A_j (e_j^*)^3} = \frac{1}{A_i (e_i^*)^3}$$

Because the pseudo-activities are strictly positive, we get:

$$\forall j, \ \forall i, \quad e_j^* = \frac{A_i^{1/3} e_i^*}{A_j^{1/3}}$$

Because $\sum_{j=1}^{n} e_j^* = 1$, we have:

$$\sum_{j=1}^{n} e_j^* = \sum_{j=1}^{n} \frac{A_i^{1/3} e_i^*}{A_j^{1/3}} = A_i^{1/3} e_i^* \sum_{j=1}^{n} \frac{1}{A_j^{1/3}} = 1$$

This leads to the expression of the equilibrium of the relative enzyme concentrations:

$$\boxed{\forall i, \quad e_i^* = \frac{A_i^{-1/3}}{\sum_{j=1}^{n} A_j^{-1/3}}} \tag{S16}$$

Introducing $e_i^*$ in equation S12, we get the flux response coefficient at equilibrium of relative enzyme concentrations:

$$R_{E_i}^{J*} = \frac{\frac{1}{A_i e_i^*}}{\sum_{k=1}^{n} \frac{1}{A_k e_k^*}}$$

and after rearrangements, we obtain:

$$\boxed{\forall i, R_{E_i}^{J*} = \frac{A_i^{-2/3}}{\sum_{j=1}^{n} A_j^{-2/3}}} \tag{S17}$$

##### SI.B.5 Evolutionary equilibrium in the case of competition (case 2)

When there is competition, $E_{\text{tot}}$ is constant and the redistribution coefficient is: $\forall i,\ \alpha_{ii} = 1$ and $\forall j \neq i,\ \alpha_{ij} = \dfrac{-e_j}{1 - e_i}$. The response coefficient becomes:

$$\forall i \quad R_{E_i}^{J} = \frac{\left( \frac{1}{A_i e_i} - \frac{e_i}{1-e_i} \sum_{j=1, j \neq i}^{n} \frac{1}{A_j e_j} \right)}{\sum_{j=1}^{n} \frac{1}{A_j e_j}} \tag{S18}$$

and the differential equation system describing the evolution of enzyme concentrations is:

$$\forall j, \quad \frac{\partial E_j}{\partial t} = 2\mu \sum_{i=1}^{n} s_i \alpha_{ij} \delta_i$$

As $E_{\text{tot}}$ is constant, this system writes for the relative concentrations:

$$\forall j, \quad \frac{\partial e_j}{\partial t} = \frac{\partial (E_j / E_{\text{tot}})}{\partial t} = \frac{1}{E_{\text{tot}}} \frac{\partial E_j}{\partial t}$$

Thus:

$$\forall j, \quad \frac{\partial e_j}{\partial t} = \frac{2\mu}{E_{\text{tot}}} \sum_{i=1}^{n} s_i \alpha_{ij} \delta_i$$

Given the relation between selection and response coefficients (equation S14), we have:

$$\forall j, \quad \frac{\partial e_j}{\partial t} \approx \frac{2\mu}{E_{\text{tot}}} \sum_{i=1}^{n} R^J_{E_i} \alpha_{ij} \frac{\delta_i^2}{E_i}$$

In addition to the purposeless solution $\forall i, \delta_i = 0$, there is an obvious solution for the steady state of relative enzyme concentrations $\frac{\partial e_i}{\partial t} = 0$:

$$\boxed{\forall i, \quad R^J_{E_i} = 0} \tag{S19}$$

From equation S18, it comes:

$$\frac{1}{A_i (e_i^*)^2} = \frac{1}{1 - e_i^*} \sum_{\substack{k=1 \\ k \neq i}}^{n} \frac{1}{A_k e_k^*}$$

Adding the case where $k = i$ in the sum, we have:

$$\frac{1}{A_i (e_i^*)^2} + \frac{1}{(1 - e_i^*) A_i e_i^*} = \frac{1}{1 - e_i^*} \sum_{k=1}^{n} \frac{1}{A_k e_k^*}$$

After rearrangements, we get:

$$\frac{1}{A_i (e_i^*)^2} = \sum_{k=1}^{n} \frac{1}{A_k e_k^*}$$

As this equality is valid for any enzyme, we have:

$$\forall i, \ \forall j \quad \sum_{k=1}^{n} \frac{1}{A_k e_k^*} = \frac{1}{A_i (e_i^*)^2} = \frac{1}{A_j (e_j^*)^2}$$

Because the pseudo-activities are strictly positive, we get:

$$e_j^* = \frac{A_i^{1/2} e_i^*}{A_j^{1/2}}$$

and because $\sum\limits_{j=1}^{n} e_j^* = 1$, we write:

$$\sum_{j=1}^{n} e_j^* = \sum_{j=1}^{n} \frac{A_i^{1/2} e_i^*}{A_j^{1/2}} = A_i^{1/2} e_i^* \sum_{j=1}^{n} \frac{1}{A_j^{1/2}} = 1$$

This leads to the equilibrium of relative concentrations when there is competition:

$$\boxed{\forall i, \quad e_i^* = \frac{A_i^{-1/2}}{\sum\limits_{j=1}^{n} A_j^{-1/2}}} \tag{S20}$$

With this expression, we can find the maximum flux value at the equilibrium:

$$J_{\text{max}} = E^0_{tot} \frac{X}{\sum\limits_{i=1}^{n} \frac{1}{A_i e_i^*}}$$

Introducing the expression of $e_i^*$ (equation S20), we get:

$$J_{\max} = E^0_{tot} \frac{X}{\sum\limits_{i=1}^{n} \frac{1}{A_i \left( \frac{A_i^{-1/2}}{\sum\limits_{j=1}^{n} A_j^{-1/2}} \right)}}$$

$$= E^0_{tot} \frac{X}{\sum\limits_{j=1}^{n} A_j^{-1/2} \sum\limits_{i=1}^{n} A_i^{-1/2}}$$

Hence:

$$\boxed{J_{\max} = E^0_{\text{tot}} \frac{X}{\left( \sum\limits_{j=1}^{n} A_j^{-1/2} \right)^2}} \tag{S21}$$

##### SI.B.6 Evolutionary equilibria in the case of co-regulation (case 3)

###### SI.B.6.1 Theoretical equilibrium

**SI.B.6.1.1 Expression of the theoretical equilibrium** When there is co-regulation between all enzymes without competition ($\forall (i,j)\ \alpha_{ij} = \beta_{ij} \neq 0$), the differential equation system becomes:

$$\frac{\partial E_j}{\partial t} = 2\mu \sum_{i=1}^{n} s_i \beta_{ij} \delta_i$$

In terms of *relative* concentrations, it writes:

$$\frac{\partial e_j}{\partial t} = \frac{\partial (E_j / E_{tot})}{\partial t} = \frac{\frac{\partial E_j}{\partial t} E_{tot} - E_j \frac{\partial E_{tot}}{\partial t}}{(E_{tot})^2} \tag{S22}$$

Because

$$\frac{\partial E_{\text{tot}}}{\partial t} = \sum_{j=1}^{n} \frac{\partial E_j}{\partial t} = \sum_{j=1}^{n} 2\mu \sum_{i=1}^{n} s_i \beta_{ij} \delta_i$$

$$= 2\mu \sum_{i=1}^{n} s_i \delta_i \sum_{j=1}^{n} \beta_{ij}$$

$$= 2\mu \sum_{i=1}^{n} s_i \delta_i B_i$$

we can write:

$$\begin{aligned}\frac{\partial e_j}{\partial t} &= \frac{E_{tot} 2\mu \sum\limits_{i=1}^{n} s_i \beta_{ij} \delta_i - E_j 2\mu \sum\limits_{i=1}^{n} s_i \delta_i B_i}{(E_{tot})^2} \\ &= \frac{2\mu}{E_{tot}} \sum_{i=1}^{n} s_i \beta_{ij} \delta_i - \frac{2\mu}{E_{tot}} e_j \sum_{i=1}^{n} s_i \delta_i B_i \\ &= \frac{2\mu}{E_{tot}} \sum_{i=1}^{n} s_i \delta_i (\beta_{ij} - e_j B_i)\end{aligned}$$

Using the expressions of $s_i$ (equation S14) and $\beta_{ij}$ (equation S2), we can write:

$$\frac{\partial e_j}{\partial t} = \frac{2\mu}{E_{tot}} \sum_{i=1}^{n} R^J_{E_i} \frac{\delta_i^2}{E_i} \left( \frac{B_i}{B_j} - e_j B_i \right)$$

and after rearrangements, we get:

$$\frac{\partial e_j}{\partial t} = \frac{2\mu}{E_{tot}^2} (1/B_j - e_j) \sum_{i=1}^{n} \frac{R^J_{E_i} \delta_i^2}{e_i} B_i \tag{S23}$$

An obvious solution for $\frac{\partial e_j}{\partial t} = 0$ for all $j$ is:

$$\boxed{\forall j, \quad e_j^* = \frac{1}{B_j}} \tag{S24}$$

Note that if at least one co-regulation coefficient is negative, leading to at least one negative $B_i < 0$ (equation S2), the equilibrium concentration is negative, which is not biologically possible.

**SI.B.6.1.2 Stability of the theoretical equilibrium** Consider a transition from a theoretical equilibrium $\mathbf{E^{t_1}}$ to a new state $\mathbf{E^{t_2}}$ due to a mutation of effect $\nu$ targeting enzyme $i$. The relative concentration of any enzyme $j \neq i$ after mutation is:

$$e_j^{t_2} = \frac{E_j^{t_2}}{E_{tot}^{t_2}} = \frac{E_j^{t_1} + \beta_{ij}\nu}{E_{tot}^{t_1} + B_i \nu}$$

Since the system is at the theoretical equilibrium at $t_1$, we have $e_j^{t_1} = e_j^* = 1/B_j$, so:

$$e_j^{t_2} = \frac{\frac{E_{tot}^{t_1}}{B_j} + \beta_{ij}\nu}{E_{tot}^{t_1} + B_i \nu}$$

and from $\beta_{ij} = B_i/B_j$ (equation S2) we obtain:

$$\begin{aligned}e_j^{t_2} &= \frac{\frac{E_{tot}^{t_1}}{B_j} + \frac{B_i}{B_j}\nu}{E_{tot}^{t_1} + B_i \nu} \\ &= \frac{1}{B_j}\end{aligned}$$

Similarly, we have for the target enzyme $i$:

$$e_i^{t_2} = \frac{E_i^{t_2}}{E_{tot}^{t_2}} = \frac{E_i^{t_1} + \nu}{E_{tot}^{t_1} + B_i\nu}$$

Because $E_i = e_i E_{\text{tot}}$ and $e_i^{t_1} = e_i^* = 1/B_i$, we have:

$$\begin{aligned} e_i^{t_2} &= \frac{\frac{E_{tot}^{t_1}}{B_i} + \nu}{E_{tot}^{t_1} + B_i\nu} \\ &= \frac{1}{B_i} \end{aligned}$$

Thus, no mutation, whatever its effect, can modify the *relative* enzyme concentrations at the theoretical equilibrium.

**SI.B.6.1.3 Response coefficient at the theoretical equilibrium** From the general expression of the flux response coefficient (equation S12), it is possible to find the expression of the response coefficient at the equilibrium when there is co-regulation. Knowing that $\alpha_{ij} = \beta_{ij}$ in the case of co-regulation, we have for replacing $e_i$ by $e_i^* = \frac{1}{B_i}$ and using the relation $B_i = \beta_{ij} B_j$ (equation S2), we get:

$$R_{E_i}^{J*} = e_i^* \frac{\sum_{j=1}^{n} \frac{\beta_{ij}}{A_j (e_j^*)^2}}{\sum_{j=1}^{n} \frac{1}{A_j e_j^*}} = \frac{1}{B_i} \frac{\sum_{j=1}^{n} \frac{B_i}{B_j} \frac{1}{A_j (1/B_j)^2}}{\sum_{j=1}^{n} \frac{1}{A_j/B_j}}$$

After rearrangements, we obtain:

$$\boxed{\forall i, R_{E_i}^{J} = 1} \tag{S25}$$

Thus, the flux $J^*$ at the theoretical equilibrium is proportional to each $E_i$. Its value is:

$$J^* = E_{tot} \frac{X}{\sum_{i=1}^{n} \frac{B_i}{A_i}}$$

**SI.B.6.2 The parametric equation of the line relating the relative enzyme concentrations.** Enzyme concentrations are all linearly related, as are the relative enzyme concentrations. So they move on a straight line of the $n$-dimensional space where each axis corresponds to the relative concentration of an enzyme.

This line, $\mathcal{E}$, has two known points: the initial point $\mathbf{e^0}$, named $\mathbf{O}$, and the theoretical equilibrium point $\mathbf{e}^*$, named $\mathbf{F}$. Thus any point $\mathbf{M}$ of coordinates $\mathbf{e}$ is on the line $\mathcal{E}$ if and only if there is a real $\tau$ such as $\overrightarrow{OM} = \tau \times \overrightarrow{OF}$, *i.e.* if:

$$\forall i, \quad e_i - e_i^0 = \tau(e_i^* - e_i^0)$$

We know that $\forall i, e_i^* = 1/B_i$, so the parametric equation of the line $\mathcal{E}$ is:

$$\boxed{\exists \tau \in \mathbb{R} \text{ such as } \forall i,\ e_i = \tau \left( \frac{1}{B_i} - e_i^0 \right) + e_i^0} \tag{S26}$$

When $\tau = 0$, $e_i = e_i^0$ and when $\tau = 1$, $e_i = 1/B_i$, which is the theoretical equilibrium.

**SI.B.6.3 Effective equilibrium in the case of negative co-regulation** In Supporting information SI.B.6.1.1, we established the differential equation system describing the evolution of relative enzyme concentrations when there is co-regulation (equation S23):

$$\forall_j, \quad \frac{\partial e_j}{\partial t} = \frac{2\mu}{E_{tot}^2}(1/B_j - e_j)\sum_{i=1}^{n} \frac{R_{E_i}^J \delta_i^2}{e_i} B_i \tag{S27}$$

This equation is cancelled if $\forall j$, $e_j = 1/B_j$, which led to the theoretical equilibrium. Actually there is another trivial solution, which leads to the effective equilibrium:

$$\boxed{\forall i, \quad R_{E_i}^J = 0} \tag{S28}$$

or, from equation S12:

$$\forall i \quad \sum_{j=1}^{n} \frac{\beta_{ij}}{A_j(\tilde{e_j})^2} = 0$$

This equality can only be true if at least one co-regulation coefficient is negative. Because $\forall (i,j), \beta_{ij} = B_i/B_j$ (equation S2), we get:

$$\forall i \quad B_i \sum_{j=1}^{n} \frac{1}{B_j A_j(\tilde{e_j})^2} = 0$$

or:

$$\boxed{\sum_{j=1}^{n} \frac{1}{B_j A_j(\tilde{e_j})^2} = 0} \tag{S29}$$

Thanks to the relationship between $e_j$ and $\tau$ (equation S26), searching for $\tilde{e}_j$ for all $j$ is equivalent to searching for $\tilde{\tau}$ using the following equation:

$$\sum_{j=1}^{n} \frac{1}{B_j A_j(\tilde{\tau}(1/B_j - e_j^0) + e_j^0)^2} = 0 \tag{S30}$$

Solving this equation for $\tilde{\tau}$ allows us to calculate the relative concentrations $\tilde{\mathbf{e}}$ at effective equilibrium.

#### SI.B.7 Evolutionary equilibrium in the case of competition plus co-regulation (case 4)

##### SI.B.7.1 Theoretical equilibrium

**SI.B.7.1.1 Expression of the theoretical equilibrium** When there is co-regulation and competition, we have $\forall i \alpha_{ii} = 1$ and $\forall (i, j \neq i)\ \alpha_{ij} = \frac{\beta_{ij} - B_i e_j}{1 - B_i e_i}$. As all enzyme are co-regulated, *i.e.* $\beta_{ij} \neq 0$ for all $(i,j)$, we can apply equation S2 to the redistribution coefficient, leading to:

$$\alpha_{ij} = \frac{1/B_j - e_j}{1/B_i - e_i} \tag{S31}$$

Note that this expression is also valid when $j = i$: $\alpha_{ii} = \frac{1/B_i - e_i}{1/B_i - e_i} = 1$. So the differential equation system becomes:

$$\forall j, \quad \frac{\partial E_j}{\partial t} = 2\mu \sum_{i=1}^{n} s_i \frac{1/B_j - e_j}{1/B_i - e_i}\delta_i = 2\mu(1/B_j - e_j)\sum_{i=1}^{n} \frac{1}{1/B_i - e_i} s_i \delta_i$$

In terms of relative enzyme concentration, we have:

$$\frac{\partial e_j}{\partial t} = \frac{\partial (E_j / E_{\text{tot}})}{\partial t}$$

or, because $E_{\text{tot}}$ is constant:

$$\frac{\partial e_j}{\partial t} = \frac{1}{E_{\text{tot}}} \frac{\partial E_j}{\partial t}$$

Hence, the differential equation system for the relative enzyme concentrations is:

$$\forall j, \quad \frac{\partial e_j}{\partial t} = 2\mu \frac{(1/B_j - e_j)}{E_{\text{tot}}} \sum_{i=1}^{n} \frac{1}{1/B_i - e_i} s_i \delta_i \tag{S32}$$

An obvious solution for the evolutionary equilibrium $\forall j \; \frac{\partial e_j}{\partial t} = 0$ is:

$$\boxed{\forall j, \quad e_j^* = \frac{1}{B_j}} \tag{S33}$$

At this point, the flux value is:

$$\boxed{J^* = E_{\text{tot}}^0 \frac{X}{\sum_{j=1}^{n} \frac{B_j}{A_j}}} \tag{S34}$$

**SI.B.7.1.2 Stability of the theoretical equilibrium.** Consider a system at theoretical equilibrium $\mathbf{E}^{\mathbf{t_1}}$ passing to another state $\mathbf{E}^{\mathbf{t_2}}$ by a mutation of effect $\nu$ affecting enzyme $i$. So concentration of any enzyme $j$ at state after mutation can be written:

$$E_j^{t_2} = E_j^{t_1} + \frac{1 - \beta_{ij} e_j}{1 - B_i e_i} \delta_i$$

And for the targeted enzyme $i$:

$$E_i^{t_2} = E_i^{t_1} + \delta_i$$

where the actual effect $\delta_i$ of the mutation is (equation S8):

$$\delta_i = \frac{1 - B_i e_i^{t_1}}{\frac{1}{\nu} + \frac{1}{E_{\text{tot}}}}$$

But because at time $t_1$, the system is at theoretical equilibrium, we have $e_i^{t_1} = \frac{1}{B_i}$, and therefore:

$$\delta_i = 0$$

So we obtain $E_j^{t_2} = E_j^{t_1}$ and $E_i^{t_2} = E_i^{t_1}$ whatever the canonical effect $\nu$ of the mutation. The system is blocked at theoretical equilibrium.

**SI.B.7.2 Effective equilibrium** In Supporting Information SI.B.7.1.1, we established the differential equation system when there is competition plus co-regulation (equation S32).

Using relationship between response and selection coefficients (equation 15), equation S32 becomes:

$$\forall j, \quad \frac{\partial e_j}{\partial t} = 2\mu \frac{(1/B_j - e_j)}{E_{\text{tot}}} \sum_{i=1}^{n} \frac{1}{1/B_i - e_i} R^J_{E_i} \frac{\delta_i^2}{E_i} \tag{S35}$$

In addition to $e_j = 1/B_j$, which is the theoretical equilibrium, the system admits another obvious solution:

$$\boxed{\forall i, \quad R^J_{E_i} = 0} \tag{S36}$$

Introducing the expression of $\alpha_{ij}$ (equation S31) in the equation of the flux response coefficient (equation S12) leads to:

$$R^J_{E_i} = \frac{e_i}{1/B_i - e_i} \frac{\sum_{j=1}^{n} \frac{1/B_j - e_j}{A_j(e_j)^2}}{\sum_{j=1}^{n} \frac{1}{A_j e_j}}$$

So canceling equation S36 amounts to:

$$\sum_{j=1}^{n} \frac{\frac{1}{B_j} - \tilde{e}_j}{A_j(\tilde{e}_j)^2} = 0$$

Thanks to the relationship between $e_j$ and $\tau$ (equation 28), we know that searching for the $\tilde{e}_j$'s that cancel this equation is equivalent to searching for a particular $\tilde{\tau}$ value. So we write:

$$\begin{aligned} \sum_{j=1}^{n} \frac{\frac{1}{B_j} - \tilde{e}_j}{A_j(\tilde{e}_j)^2} &= \sum_{j=1}^{n} \frac{1/B_j - (\tau(1/B_j - e_j^0) + e_j^0)}{A_j(\tilde{\tau}(1/B_j - e_j^0) + e_j^0)^2} \\ &= (1-\tau) \sum_{j=1}^{n} \frac{(1/B_j - e_j^0)}{A_j(\tilde{\tau}(1/B_j - e_j^0) + e_j^0)^2} \end{aligned}$$

In addition to the trivial solution $\tau = 1$, which corresponds to the theoretical equilibrium, the other solution is the $\tau$ value that cancels this equation:

$$\boxed{\sum_{j=1}^{n} \frac{1/B_j - e_j^0}{A_j(\tilde{\tau}(1/B_j - e_j^0) + e_j^0)^2} = 0} \tag{S37}$$

Once $\tilde{\tau}$ is found, it is easy to find the relative concentrations $\tilde{\mathbf{e}}$ at effective equilibrium.

##### SI.B.8 Expressing absolute enzyme concentrations in function of the driving variable

**SI.B.8.1 In the case of co-regulation** In the parametric equation S26, we can make visible the *absolute* enzyme concentrations, using the equation $e_i = E_i/E_{\text{tot}}$:

$$\forall i, \frac{E_i}{E_{\text{tot}}} = \tau \left( \frac{1}{B_i} - \frac{E_i^0}{E_{\text{tot}}^0} \right) + \frac{E_i^0}{E_{\text{tot}}^0} \tag{S38}$$

As there is not competition, $E_{\text{tot}}$ varies with $\tau$. The relation between $E_{\text{tot}}$ and $\tau$ can be derived from the relation $B_i = \frac{\Delta E_{\text{tot}}}{\Delta E_i}$ (equation 6). We have:

$$\begin{aligned} E_{\text{tot}} &= E^0_{\text{tot}} + B_i(E_i - E^0_i) \\ &= E_{\text{tot}} B_i e_i + E^0_{\text{tot}}(1 - B_i e^0_i) \\ &= E^0_{tot} \frac{1 - B_i e^0_i}{1 - B_i e_i} \end{aligned}$$

Replacing $e_i$ by its expression (equation S26), we get:

$$E_{tot} = E^0_{tot} \frac{1 - B_i e^0_i}{1 - B_i(\tau(1/B_i - e^0_i) + e^0_i)}$$

which is simplified to (for $\tau \neq 1$):

$$E_{\text{tot}} = \frac{E^0_{\text{tot}}}{1 - \tau} \tag{S39}$$

Introducing this expression of $E_{\text{tot}}$ in equation S38, we get:

$$\forall i, \frac{E_i(1 - \tau)}{E^0_{\text{tot}}} = \tau \left( \frac{1}{B_i} - \frac{E^0_i}{E^0_{\text{tot}}} \right) + \frac{E^0_i}{E^0_{\text{tot}}} \tag{S40}$$

Thus:

$$E_i = E^0_i + \frac{\tau E^0_{\text{tot}}}{(1 - \tau) B_i} \tag{S41}$$

Or:

$$\boxed{E_i = E^0_i + \frac{E^0_{\text{tot}}}{B_i} \left( -1 + \frac{1}{1 - \tau} \right)} \tag{S42}$$

**SI.B.8.2 In the case of co-regulation and competition** When there is both competition and co-regulation, the difference with the previous case is that $E_{\text{tot}}$ is constant. So, replacing $e_i$ by $E_i/E_{\text{tot}}$ in equation S26, we get directly:

$$\boxed{E_i = \tau \left( \frac{E_{\text{tot}}}{B_i} - E^0_i \right) + E^0_i} \tag{S43}$$

##### SI.B.9 Evolution model of the driving variable

We assume here that the mutations target a factor controlling the concentrations of a group of enzymes. Formally, this corresponds to mutations affecting the driving variable $\tau$:

$$\tau^m = \tau^r + \nu_\tau$$

where $\nu_\tau$ is the effect of the mutation on $\tau$.

**SI.B.9.1 Redistribution coefficient and mutation effect** If the mutation affects $\tau$, the questions arise of the actual effect $\delta_i$ of this mutation on a particular enzyme $i$ and of the fate of the redistribution coefficients $\alpha_{ij}$. We examine these questions in the two possible situations.

**SI.B.9.1.1 In the case of co-regulation** For all enzyme $i$, we have from equation S42:

$$E_i^r = E_i^0 + \frac{E_{\text{tot}}^0}{B_i}\left(-1 + \frac{1}{1-\tau^r}\right)$$
$$\text{and } E_i^m = E_i^0 + \frac{E_{\text{tot}}^0}{B_i}\left(-1 + \frac{1}{1-\tau^r-\nu_\tau}\right)$$

leading to the actual effect of the mutation on enzyme $i$:

$$\delta_i = E_i^m - E_i^r = \frac{E_{\text{tot}}^0}{B_i}\left(\frac{1}{1-\tau^r-\nu_\tau} - \frac{1}{1-\tau^r}\right) \tag{S44}$$

So $\delta_i$ depends both on $\nu_\tau$ and $B_i$.

For two enzymes $i$ and $j$, we have from the previous equation:

$$(E_i^m - E_i^r)\frac{B_i}{E_{\text{tot}}^0} = (E_j^m - E_j^r)\frac{B_j}{E_{\text{tot}}^0} = \left(\frac{1}{1-\tau^r-\nu_\tau} - \frac{1}{1-\tau^r}\right)$$

Therefore, we obtain:

$$E_j^m = E_j^r + \frac{B_i}{B_j}(E_i^m - E_i^r)$$

and because $B_i/B_j = \beta_{ij}$, we have:

$$E_j^m = E_j^r + \beta_{ij}(E_i^m - E_i^r) \tag{S45}$$

Thus, the equation $\Delta E_j = \beta_{ij}\Delta E_i$ between two co-regulated enzymes is valid, whether enzyme $i$ or the driving variable is the target of the mutation.

**SI.B.9.1.2 In the case of competition and co-regulation** For all enzyme $i$, we have from equation S43:

$$E_i^r = E_i^0 + \tau^r\left(\frac{E_{\text{tot}}^0}{B_i} - E_i^0\right)$$
$$\text{and } E_i^m = E_i^0 + (\tau^r + \nu_\tau)\left(\frac{E_{\text{tot}}^0}{B_i} - E_i^0\right)$$

leading to the actual effect of the mutation on enzyme $i$:

$$\delta_i = E_i^m - E_i^r = \nu_\tau\left(\frac{E_{\text{tot}}^0}{B_i} - E_i^0\right) \tag{S46}$$

Again $\delta_i$ depends both on $\nu_\tau$ and $B_i$.

For two enzymes $i$ and $j$, we have from equation S46:

$$\frac{E_i^m - E_i^r}{\frac{E_{\text{tot}}^0}{B_i} - E_i^0} = \frac{E_j^m - E_j^r}{\frac{E_{\text{tot}}^0}{B_j} - E_i^0} = \nu_\tau$$

Therefore, we obtain:

$$E_j^m = E_j^r + \frac{E_{\text{tot}}^0/B_j - E_j^0}{E_{\text{tot}}^0/B_i - E_i^0}(E_i^m - E_i^r)$$

and because $\frac{E_{\text{tot}}^0/B_j - E_j^0}{E_{\text{tot}}^0/B_i - E_i^0} = \alpha_{ij}$ for two co-regulated enzymes (Supporting Information SI.B.1.4), we have:

$$E_j^m = E_j^r + \alpha_{ij}(E_i^m - E_i^r) \tag{S47}$$

Thus, the equation $\Delta E_j = \alpha_{ij}\Delta E_i$ between two co-regulated enzymes is valid, whether enzyme $i$ or the driving variable is the target of the mutation.

###### SI.B.9.2 Evolution model of the driving variable and evolutionary equilibria

With the same assumptions as for the evolution model of enzyme concentrations, the variation of the driving variable $\tau$ can be represented as a stochastic process. At each time $t$, its rate of variation can be described with the following complete system of events:

$$\left(\frac{\partial\tau}{\partial t}\right)^{(t)} = \begin{cases} \textbf{Probability} & \textbf{Mutation effect} & \textbf{Event} \\ \mu_\tau 2s_\tau^{(t)} & \nu_\tau^{(t)} & \text{Fixation of mutation targeting } \tau \\ 1-\mu_\tau 2s_\tau^{(t)} & 0 & \text{No mutation or no fixation} \end{cases} \quad \text{(S48)}$$

where $\mu_\tau$ is the mutation rate and $s_\tau$ the coefficient of selection ($s_\tau = \dfrac{J_\tau^m - J^r}{J^r}$).

At each time $t$ the *average* variation of the driving variable $\tau$ is:

$$\frac{\partial\tau}{\partial t} = \mu_\tau 2 s_\tau \nu_\tau \quad \text{(S49)}$$

Assuming that mutation effects $\nu_\tau$ are small compared to $\tau$, we have (Supporting Information SI.B.3.1) :

$$s_\tau = \left(\frac{\Delta J}{J}\right)_\tau = \frac{\Delta J}{\Delta\tau}\frac{\Delta\tau}{J} \approx \frac{\partial J}{\partial\tau}\frac{\nu_\tau}{J}$$

So equation S49 can be rewritten as:

$$\frac{\partial\tau}{\partial t} \approx 2\mu_\tau \frac{\partial J}{\partial\tau}\frac{\nu_\tau^2}{J} \quad \text{(S50)}$$

To find the evolutionary equilibrium of the driving variable $\tau$, we have to search for $\tau$ such as $\dfrac{\partial\tau}{\partial t} = 0$, which is equivalent, from the previous equation, to find a solution for $\dfrac{\partial J}{\partial\tau} = 0$. We can decompose this equation:

$$\begin{aligned} \frac{\partial J}{\partial\tau} &= \sum_{i=1}^{n} \frac{\partial J}{\partial E_i}\frac{\partial E_i}{\partial\tau} \\ &= \sum_{i=1}^{n} R_{E_i}^J \frac{J}{E_i}\frac{\partial E_i}{\partial\tau} \end{aligned}$$

So canceling this equation amounts to canceling the flux response coefficients $R_{E_i}^J$ for all $i$, which is the definition of the effective equilibrium of relative concentrations. From that, we can find $\tilde{\tau}$ in the cases of negative co-regulation (Supporting Information SI.B.6.3) and competition plus co-regulation (Supporting Information SI.B.7.2).

Regarding the theoretical equilibrium in the case of co-regulation plus competition, cancelling the derivative of equation S43

$$\frac{\partial E_i}{\partial\tau} = \frac{E_{\text{tot}}}{B_i} - E_i$$

leads to $e_i^* = 1/B_i$, which is the theoretical equilibrium. In the case of positive co-regulation alone, $J$ increases continuously. As a consequence, $\dfrac{\delta\tau}{\delta t}$ tends towards 0 (see equation S50), which corresponds to $\tau = 1$, the theoretical equilibrium (Supporting information SI.B.6.2).

To conclude, in the case of co-regulation, the evolution model is modified if the mutations target the driving variable $\tau$ rather than concentrations of individual enzymes, but the evolutionary equilibria (theoretical and effective) do not change.
